# Plasminogen activator inhibitors orchestrate the immunosuppressive tumor microenvironment in pancreatic cancer

**DOI:** 10.1101/2025.06.11.659098

**Authors:** Chiara Falcomatà, Sebastian R. Nielsen, Maximilian M. Schaefer, Bhavya Singh, Alexander Tepper, Divya Chhamalwan, Hunter T. Potak, Maxime Dhainaut, Gurkan Mollaoglu, Matthew D. Park, Miriam Merad, Alessia Baccarini, Brian D. Brown

## Abstract

Pancreatic ductal adenocarcinoma (PDAC) is characterized by a dense extracellular matrix (ECM) that sustains an immunosuppressive tumor microenvironment (TME). While this protective niche has been described, the molecular determinants orchestrating its formation and dictating its immune interactions are not well defined. Using Perturb-map, we determine how dozens of different gene perturbations shape the growth and cellular environments of PDAC clones through space and time. Our study reveals dynamic, gene-specific adaptations of immune neighborhoods during clonal selection. We identified *Serpinb2* (PAI2) and *Serpine1* (PAI1) as key cancer-derived mediators of TME remodeling and immune evasion. These factors promote the deposition of a fibrin-rich ECM that shapes immune cell composition, locally retains and polarizes immunosuppressive macrophages and excludes cytotoxic T cells. Deletion of either *Serpinb2* or *Serpine1* greatly enhanced tumor response to anti-PD1 immunotherapy in an aggressive PDAC model. Transcriptomic analysis further linked their expression to distinct PDAC subtypes and poor patient survival. Our findings demonstrate that *Serpinb2* and *Serpine1* establish a permissive niche for tumor progression and show how PDAC cells exploit components of the fibrinolysis pathway to remodel the ECM, alter macrophage composition, and protect themselves from immune editing, ultimately reinforcing the role of extracellular factors in shaping an immune-privileged tumor niche.

## Introduction

Pancreatic ductal adenocarcinoma (PDAC) is one of the most lethal malignancies, with a five-year survival rate of only 12% and a median survival of less than six months^1^. It is now the third leading cause of cancer-related mortality^1^, reflecting its exceptional therapeutic resistance and the urgent need for new treatment strategies. While immunotherapies have improved the treatment and prognosis of several types of cancer, their use has not been effective in PDAC so far. The composition of the tumor microenvironment (TME) is considered a key factor in immunotherapy resistance, with T cell infiltration being one of the strongest correlates of immunotherapy response. The TME of PDAC is often characterized by limited CD8+ T cells infiltration. While a fraction of PDACs are almost completely devoid of CD8+ T cells, in those where T cells are present, they frequently exhibit features of exhaustion. This is believed to be due to an enrichment of immunosuppressive immune cells, non-malignant stromal cells, and to the deposition of a dense extracellular matrix (ECM), which can obstruct T cell infiltration and cytotoxic activity^2–4^. Precisely how PDAC establishes its immunosuppressive TME is still poorly understood, but mounting evidence points to the critical role of tumor-derived extracellular factors^3,5^.

Cancer cells are not isolated entities; they actively engage in dynamic crosstalk with their surroundings, orchestrating the TME through secreted factors, receptor-ligand interactions, and other extracellular signals^6^. This communication is particularly relevant in PDAC, where tumor-driven signaling cascades likely underpin the immunosuppressive and desmoplastic nature of the TME^2–4^. Advances in single-cell RNA sequencing (scRNA-seq) and spatial transcriptomics have provided insight into the spatial and molecular features of tumors, enabling the identification of receptor-ligand interactions and other extracellular mediators at single-cell resolution^7–11^. However, while these studies have cataloged potential players in shaping PDAC’s TME^2,3^, the functional contribution of many of these factors remains poorly defined. Scalable approaches aimed at dissecting the function of multiple genes *in vivo* are still limited. Classical pooled CRISPR screens, though powerful for high-throughput analyses, are often restricted to identifying cell-intrinsic processes and cannot assess genes extracellular functions, such as immune cell recruitment, changes in localization of cell types within tissues, or the functions of secreted factors. This significant gap highlights the need for systematic studies capturing the functional and spatial complexity of the PDAC TME.

Here, we sought to identify extracellular factors that promote PDAC growth and immune evasion. We curated a list of prioritized genes implicated in cancer-host interactions, leveraging insights from scRNA-seq, genetic perturbations, and existing literature. To evaluate their functional roles, we employed Perturb-map, a spatial functional genomics platform, which enables the effects of many different perturbations on the TME to be resolved *in situ* and simultaneously^12,13^. We found that several extracellular factors significantly influence tumor growth and the PDAC TME. Notably, genes such as *Pthlh* and *Ly6d*, which had minimal impact *in vitro*, profoundly affected *in vivo* tumor fitness, driving changes in immune cell recruitment and TME dynamics. Two inhibitors of the fibrinolytic pathway, *Serpinb2* (PAI2) and *Serpine1* (PAI1), emerged as central orchestrators of PDAC progression and immune suppression. Their genetic ablation altered the ECM composition of the cancer cells neighborhood and reduced local retention and programing of immunosuppressive macrophages, enabling CD8+ T cells increased access to the PDAC cells. While the role of the ECM as a physical barrier to anti-tumor immunity in PDAC is well-recognized, our findings uncover a previously unappreciated mechanism by which *Serpinb2*- and *Serpine1*-driven fibrin-rich ECM actively orchestrates immune suppression. This fibrin-rich ECM not only facilitates immunosuppressive macrophage recruitment and immune evasion but also establishes a critical therapeutic vulnerability, sensitizing tumors to PD1 checkpoint blockade. Targeting this axis could undo immunosuppressive barriers and enhance the efficacy of immunotherapies for PDAC.

## Results

### Identification and prioritization of extracellular factors shaping the PDAC TME

Owing to its importance in PDAC progression, we sought to identify genes involved in orchestrating the PDAC TME. We analyzed scRNA-seq datasets from human^14^ and mouse^15^ PDAC, focusing on genes predominantly expressed in premalignant or tumor cells (adjusted p-value ≤ 0.05, log2 fold change > 4 compared to control) (**Figure S1a**). To pinpoint genes with extracellular activity, and thus capable of intercellular interactions, we performed enrichment analysis using the Compartments database for protein subcellular localization^16^, and identified the genes falling within the enriched term “extracellular region part” across comparisons. Genes meeting this criterion were compared across differential expression analyses (**Figure S1b**) and further assessed for viability effects using DepMap data from 45 human pancreatic tumor cell lines (**Figure S1c**). We prioritized genes with minimal *in vitro* fitness effects (DepMap CRISPR scores between -0.15 and 0.15). This analysis yielded a diverse set of genes encoding cell surface and secreted proteins, including genes involved in extracellular matrix assembly and organization, immune responses, and, interestingly, coagulation pathways (**Figure 1a**). To expand functional breadth, we incorporated additional candidates based on potential redundancy within pathways, culminating in a selection of 34 genes for further investigation (**Figure 1a**, **1b**). Notably, our analysis revealed consistent upregulation of genes associated with cell-cell communication in PDAC, with gene families such as IL-1, Mucins, and Serpins prominently represented (**Figure 1b**), some of which have already been linked to malignant progression, including *Il1b*^17,18^ and *Il33*^19,20^. Competition, compensation, or hierarchical relationships among these extracellular factors could play a role in shaping the TME, influencing their relative contributions to tumor progression and immune modulation. However, the causal impact of many of them on PDAC growth and TME formation remains poorly understood.

**Figure 1.**
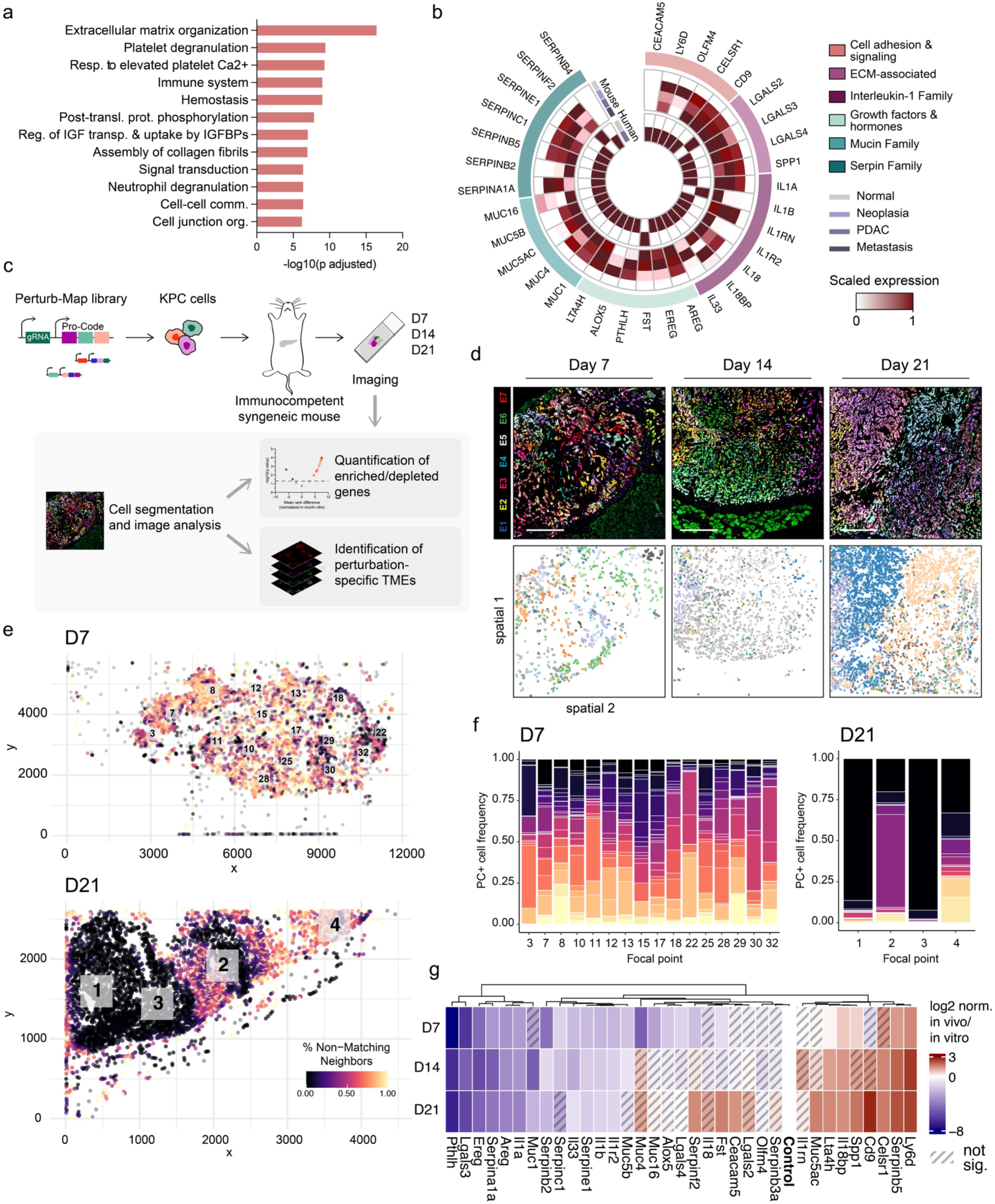
Perturb-map identifies cancer-cell derived extracellular factors regulating PDAC growth *in vivo*. **a**, Enrichment analysis of the top Reactome pathways based on 310 genes identified from the CRISPR/Cas9 dependency analysis in **Figure S1**. The top 12 pathways are displayed. **b**, Circos plot showing the scaled expression levels of prioritized genes across human and mouse scRNA-seq datasets. The outer circle categorizes genes by functional groups. **c**, KPC cells transduced with the Perturb-Map CRISPR/Cas9 library were orthotopically transplanted into syngeneic Cas9-expressing mice. Tumors were harvested at days 7, 14, and 21 post-transplantation, fixed, formalin-embedded, and stained for analysis. Tumor samples from 5, 7, and 8 mice were analyzed at each time point respectively, with two images captured per mouse. **d**, Representative images of orthotopic pancreatic tumors at various stages (35 PC, 7 tags) with corresponding digital reconstructions displayed below each image. Each color represents a distinct unique gene KO in the digital reconstruction. Scale bar: 250 μm. **e,** Representative digital reconstruction of a tumor section isolated 7 days (above) and 21 days (below) following orthotopic transplantation. The color scale indicates the percentage of non-matching neighbors, where yellow represents areas with higher heterogeneity, and black represents more homogeneous regions. White squares mark focal areas, identified using mean-shift clustering to detect regions of high local cell density based on a defined bandwidth. **f,** Frequency of PC+ cells within the focal areas identified in panel (**e**), highlighting the distribution of specific cell populations. Data for day 7 are represented on the left, day 21 on the right. Each color in the bar graph represents a distinct PC+ (i.e. KO) population of tumor cells. **g,** Log2 fold change of the ratio between normalized *in vivo* and *in vitro* viability scores for each of the indicated genes across all analyzed images at different time points, relative to the internal library control. Genes with an adjusted p value > 0.05 compared to the internal control are marked with dashed lines. Multiple Mann-Whitney tests were used to estimate significance. PDAC: pancreatic ductal adenocarcinoma, PC: pro-code, scRNA-seq: single cell RNA sequencing.

### *In situ* detection of Pro-Code labelled PDAC populations by imaging

Before applying Perturb-map to determine the functions of the prioritized genes, we sought to better understand the growth pattern of pancreatic cancer clones in a syngeneic, orthotopic *in vivo* model of PDAC. We transduced KPC (*Kras^G12D^;Trp53^R^*^172^*^H^*) PDAC cells with a library of lentiviral vectors encoding 35 unique Pro-Codes (PCs). PCs are protein-based barcodes composed of triplet combinations of linear epitopes fused to a scaffold protein, in this case nuclear mCherry (**Figure S2a**). The PC-labeled KPC cells were orthotopically implanted into the pancreas of immunocompetent mice (**Figure S2b**). Two weeks post-implantation, tumors were harvested and analyzed by multiplexed imaging^21^ to spatially resolve the specific epitopes, and thus Pro-Code, expressed by each cancer cell (**Figure S2b, S2c**).

We compared epitope frequency profiles of tumor cells cultured *in vitro* (assessed via CyTOF) with those from *in vivo* tumors post-implantation. PC frequencies were highly consistent pre- and post-implant, with a maximum *in vitro*-to-*in vivo* deviation of 4.8% (**Figure S2d**). Furthermore, analysis of normalized frequencies for each unique PC epitope in the tumors revealed no significant deviations or evidence of preferential engraftment or rejection (**Figure S2e**). These data indicate the Pro-Codes provided neutral labeling of pancreatic cancer cell clones.

We next investigated the spatial growth patterns of barcoded orthotopic KPC tumors. Using cell segmentation, assignment, and digital reconstruction, we analyzed colocalization among PC+ populations. Pairwise neighbor interactions were quantified by calculating Z-scores, comparing observed interactions to randomized cell label assignments (see Methods). This analysis revealed that KPC tumors predominantly exhibit homogeneous growth of PC+ populations within defined regions (**Figure S2f**). Homogeneous growth could indicate a reliance on cell-cell cooperation or shared resource pools within clones, as opposed to competitive interactions seen in more heterogeneous models. This is interesting, as the growth pattern differed from our previous studies with 4T1 breast cancer cells implanted in the mammary fat pad, which showed high heterogeneity, with most cancer cells neighboring a different clone rather than growing as homogeneous tumor clones^12^. This highlights how tumor cell features and tissue context influence growth dynamics, and establishes this platform as a means for scaled spatial analysis of PDAC growth *in situ*.

### Functional and spatial dynamics of candidate gene KOs in PDAC progression

To evaluate the functional relevance of the candidate genes to PDAC biology, we utilized Perturb-map^12^. A Pro-Code/CRISPR library was constructed to knock-out (KO) the 34 prioritized genes, along with an unexpressed control gene (F8) (**Figure S3a**), and the library was introduced into KPC PDAC cells (**Figure 1c and S3a**). To assess the impact of the targeted genes on cell fitness, we compared PC/CRISPR representation in KPC cells with and without Cas9 after 14 days of culture. No significant differences were observed for any PC/CRISPR construct, indicating that none of the genes affected KPC cell fitness *in vitro* (**Figure S3b**). This finding was further corroborated by DepMap data, which showed that KO of the selected 34 genes in 45 human PDAC cell lines also had no significant impact on cell growth *in vitro* (**Figure S3c**). Together, these results indicate that the targeted genes do not regulate the intrinsic fitness of PDAC cells.

Next, we orthotopically implanted the KPC^PC/CRISPR^ cell pool into syngeneic, immunocompetent mice. To capture the dynamic evolution of the tumor and its microenvironment over time, we performed a time course experiment. Pancreatic tumors were harvested on days 7, 14, and 21 post-implantation, representing early, intermediate, and late growth stages, respectively (**Figure 1c**, **S3a**). We performed multiplex imaging on tumor sections to detect the Pro-Code and markers of T cells (CD8a, CD4 and FOXP3), macrophages (F4/80), B cells (B220) and dendritic cells (CD11c). After registration segmentation, and Pro-Code assignment, we identified and distinguished all cancer cells expressing distinct PC/CRISPR and nearby immune cells (**Figure 1c**, **1d**).

Imaging of tumors over time revealed distinct spatial and clonal evolution of PC+ tumor cells (**Figure 1d**). At early time points (day 7), tumors exhibited high heterogeneity, with areas of intermixing among different gene KO cell populations (i.e. positive for a specific PC), as indicated by the color scale representing the percentage of non-matching neighbors (yellow for higher heterogeneity, black for more homogeneous regions) (**Figure 1e**). By day 21, gene KO tumor cells showed a more confined and homogeneous growth pattern, with distinct gene KO populations dominating specific regions. Quantitative analysis of high local cell density focal areas further supported this trend. At day 7, the frequency of specific gene KO cells within focal areas (**Figure 1f**, left) revealed a more even distribution across different populations, reflecting the highly intermixed state of tumor cells. By day 14, and even more by day 21, the distribution shifted towards a dominance of single populations within focal areas (**Figure S4a**, **S4b** and **1f**, right), highlighting the emergence of spatially consolidated clonal regions. Thus, the homogeneity seen at late stage does not merely reflect how the tissue is seeded but is a result of a competitive evolution in which distinct KPC^PC/CRISPR^ populations become more consolidated within specific regions as cancer clones grow and interact with the microenvironment.

Next, to assess the impact of each gene KO on tumor growth, we quantified the abundance of cancer cells expressing a specific PC/CRISPR construct. PC/CRISPR counts were normalized against the control KO, treating individual animals as biological replicates to calculate normal distributions for each KO. At day 7, most gene KOs were present at a frequency comparable or modestly reduced compared to the control gene KO. However, over time, as tumors progressed and adapted to their surrounding microenvironment, the fitness differences solidified. By late time point (day 21), 11 KOs were significantly enriched and 12 were significantly depleted compared to the control (**Figure 1g**). Genes such as *Pthlh*^22^ and *Il33*^19,20^, which are of known relevance in PDAC, were depleted. Also depleted were several members of the Serpin family - including *Serpine1* and *Serpinb2*. In contrast, genes such as *Ly6d*, *Serpinb5*, and *Celsr1*, which are less functionally characterized in PDAC, were enriched. As these genes did not directly affect the intrinsic fitness of cancer cells *in vitro*, and they operate mostly extracellularly, the influence of these genes on tumor growth *in vivo* likely reflects their roles in modulating tumor-TME interactions. Our data also suggested some genes functioning in tumor plasticity and adaptation at a late stage of growth, as cancer cells with KO of *Fst*, *Serpinf2*, and *Lta4h,* only showed significant expansion after day 14. In contrast, KOs such as *Serpinb2* and *Serpine1* exhibited strong and consistent fitness defects across time points.

### Perturb-map uncovers local remodeling of the tumor microenvironment over time

Perturb-map analysis enabled us to detect the composition and spatial positioning of key immune cell types within the tumor mass (**Figure 2a**), and we could use this information to determine how each gene influenced its local cellular environment. To do this, we performed neighborhood enrichment analysis to evaluate the relative density of specific immune cell populations in proximity to cancer cells carrying different gene KOs (**Figure 2b**). We analyzed >4.5 million cancer and immune cells in total, and this revealed major spatiotemporal differences between KO clones, with distinctions evident as early as 7 days but becoming even more pronounced by day 21 (**Figure 2c**). Remarkably, despite the inherent heterogeneity of the PDAC TME - where distinct KO cancer cells are situated side by side (**Figure 1e**) - we observed consistent and reproducible patterns of immune cell distribution, which were maintained over time (**Figure 2c**).

**Figure 2.**
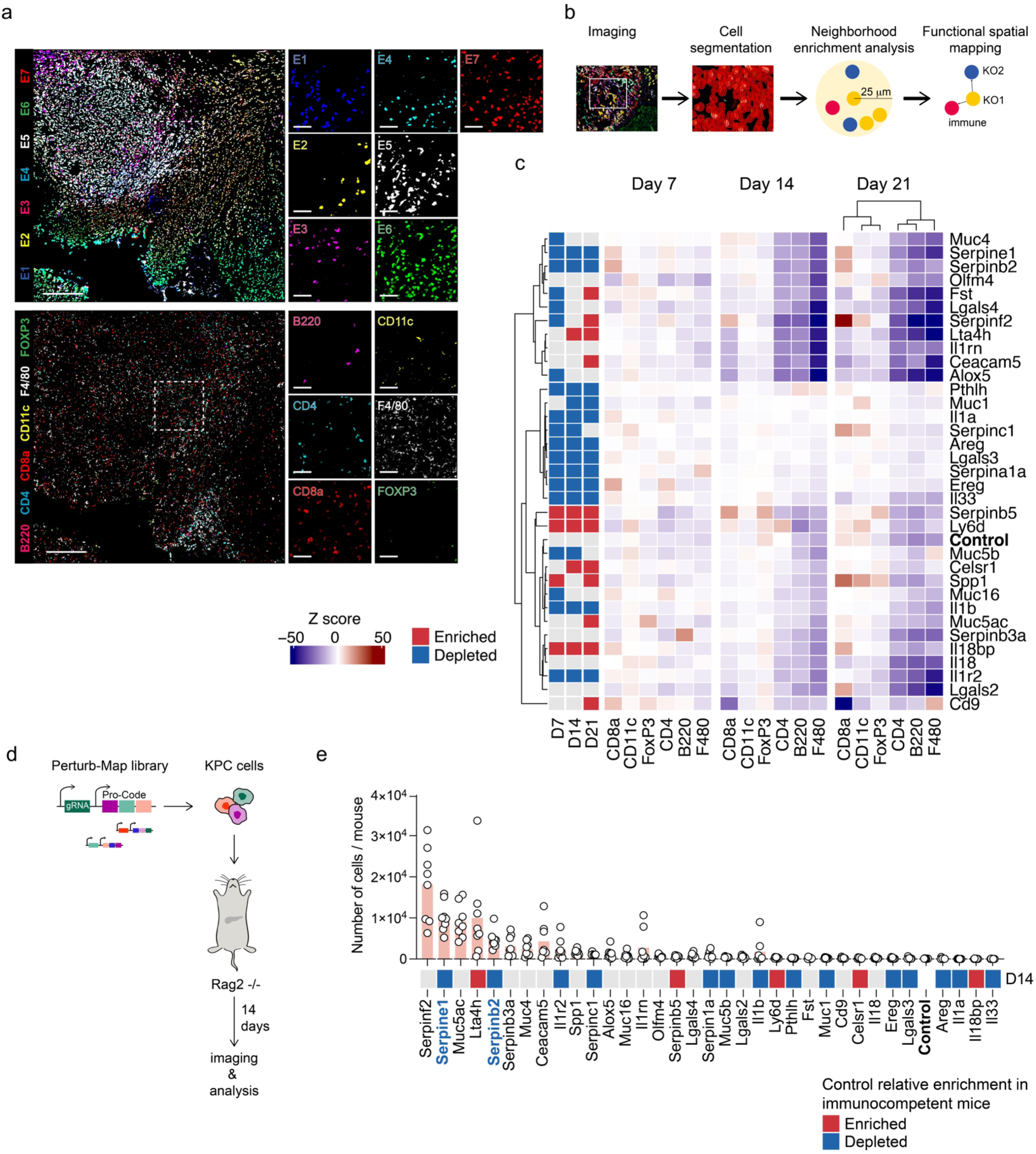
Perturb-map finds tumor-derived extracellular factors dynamically shaping the PDAC’s TME. **a,** Representative images of orthotopic pancreatic tumors stained for single PC epitopes (top, 35 PC, 7 epitope tags (E)) and immune markers (bottom). Scale bars: 250 μm for the main images and 50 μm for the insets. **b**, Schematic representation of the PDAC-TME analysis. Neighborhood enrichment analysis, using a 25 μm radius, was performed to identify immune cells surrounding each KO tumor cell. **c**, Neighborhood enrichment analysis showing immune cell enrichment within a 25 μm radius of each PDAC cell harboring a genetic KO across time points (days 7, 14, and 21). Genes are clustered based on their TMEs. The heatmap is color-coded based on the significance of the interaction relative to a permuted null distribution generated by swapping labels (1000 permutations). The relative enrichments (red) or depletions (blue), compared to the internal library control at day 7, 14 and 21, are displayed on the left. **d**, Schematic of the experimental set-up. KPC^PC/CRISPR^ cells were orthotopically transplanted into immunodeficient Rag2-/- mice. Tumors were harvested two weeks post-injection, fixed, formalin-embedded, and stained. One tumor section from 8 individual mice was analyzed. **e**, The number of KPC cells per gene KO in each mouse is shown for all genes in the Perturb-map library. Depletion or enrichment for D14, relative to the internal control in immunocompetent mice, is displayed below. *Serpinb2* and *Serpine1* are highlighted in blue. PDAC: pancreatic ductal adenocarcinoma, PC: pro-code, TME: tumor microenvironment, KO: knock-out.

Clustering of neighborhood enrichment Z-scores for each gene KO identified two distinct groups, each characterized by similar TMEs and associated with a fitness disadvantage relative to the internal control (**Figure 2c**). The first cluster, which included *Serpinb2* and *Serpine1*, displayed an increased enrichment of CD8a-positive cells, coupled with a notable decrease in CD4, B220, and F4/80-positive cells, indicative of a less immunosuppressed cellular neighborhood. In contrast, the second cluster, which encompassed nine genes, including *Pthlh*, *Muc1*, and *Il1a*, was associated with a more immune-rich environment, also showing a greater abundance of cells of the myeloid lineage (**Figure 2c**). Interestingly, loss of *Cd9* was initially neutral on clonal growth, but led to enrichment by end-stage, which was concomitant with a neighborhood enrichment in tumor macrophages and depletion of CD8 T cells. These findings provide insight into how each of the 34 cancer expressed genes influences the local immune environment of a pancreatic cancer cell. Of note, our results show that the neighborhood takes shape by day 14, when pronounced clonal heterogeneity persists. This supports the idea that specific genes regulating the proximal immune neighborhood play a key role in shaping clonal fitness and driving the development of more homogeneous clonal regions within PDAC.

### Adaptive immunity serves as a gatekeeper for *Serpinb2* and *Serpine1* tumor growth

To determine the functional contribution of the adaptive immune system in shaping the gene KO tumor phenotypes, we orthotopically transplanted the KPC^PC/CRISPR^ cell pool into the pancreas of Rag2-/- mice, which lack T cells and B cells (**Figure 2d**). In this immunodeficient context, a number of gene KOs grew more aggressively than in the immunocompetent setting (**Figures S5a-c**). Notably, genes of the Serpin family, as well as *Muc5ac* and *Lta4h*, were among the fastest growing in comparison to the control KO. Strikingly, both *Serpinb2* and *Serpine1* were amongst the most enriched in the Rag2-/- (**Figure 2e**). The difference was stark given that *Serpinb2* and *Serpine1* were highly depleted in immunocompetent context at the same time point. These findings suggest that, in the absence of an adaptive immune system, the loss of *Serpinb2* and *Serpine1* confers a growth advantage to cancer cells. Conversely, in an immunocompetent setting, their KO promotes tumor clearance. This duality is particularly notable, as the loss of *Serpinb2* and *Serpine1* is also associated with a TME enriched in CD8+ T cells and depleted in macrophages (**Figure 2c**), suggesting that their role in enhancing tumor fitness in an immunocompetent context may be related to their influence on positioning immune cells.

### PDAC derived *Serpinb2* and *Serpine1* are modulators of the tumor microenvironment

*Serpine1* encodes plasminogen activator inhibitor 1 (PAI1) and *Serpinb2* encodes PAI2. They are both serine protease inhibitors and are known to block urokinase plasminogen activator (uPA) and tissue plasminogen activator (tPA) activity^23^. Analysis of bulk RNA-seq from the TCGA and GTEx revealed that *SERPINB2* and *SERPINE1* were preferentially expressed in cancer compared to normal tissues across multiple malignancies, and especially in PDAC (**Figure 3a**).

**Figure 3.**
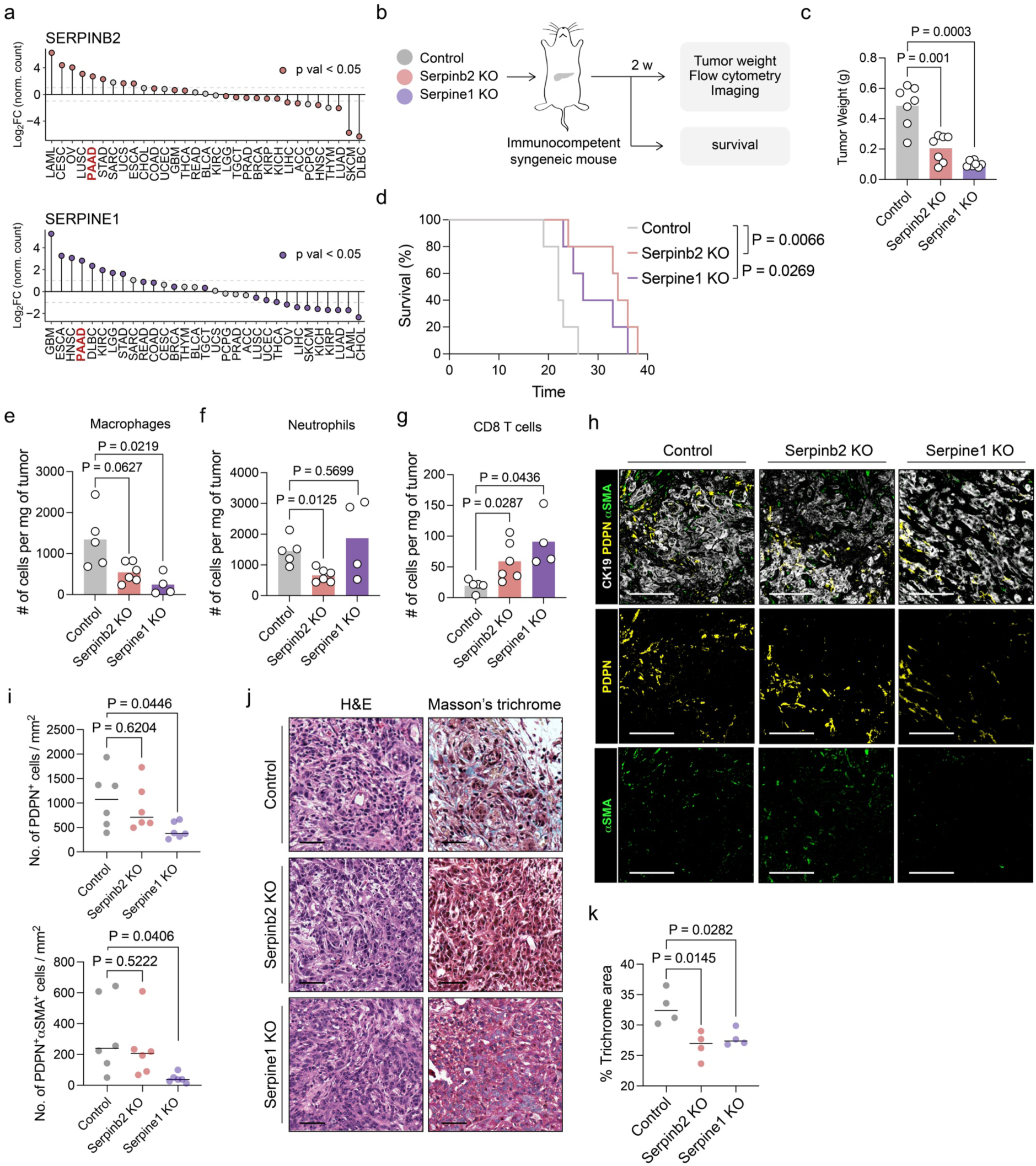
*Serpinb2* and *Serpine1* KOs exhibit anti-tumor effects and remodel the TME. **a**, Log2 fold change of bulk gene expression (TPM) for *SERPINB2* (top) and *SERPINE1* (bottom) across various human tumor types compared to normal tissues. Pancreatic cancer datasets are labeled in red. Non-significant comparisons are depicted in grey lollipops. Data were derived from TCGA and GTEx. **b**, Schematic of the experimental workflow used to validate the phenotypes following individual CRISPR/Cas9 mediated *Serpinb2* and *Serpine1* KOs. **c**, Tumor weight comparison between control, *Serpinb2* KO, and *Serpine1* KO (control n=7, *Serpinb2* KO n=7, *Serpine1* KO n=7). **d**, Kaplan–Meier survival curves of mice with control, *Serpinb2*, and *Serpine1* KOs in orthotopic transplantation models (control n=5, *Serpinb2* KO n=5, *Serpine1* KO n=5). **e-g**, Flow cytometry analysis of tumors isolated from control, *Serpinb2*, and *Serpine1* KO mice two weeks after orthotopic transplantation. The absolute number of macrophages (**e**), neutrophils (**f**), and CD8+ T cells (**g**) are shown (control n=5, *Serpinb2* KO n=6, *Serpine1* KO n=4). **h**, Pseudocolored immunohistochemistry of tumors from control, *Serpinb2*, and *Serpine1* KOs, isolated two weeks post-transplantation. Sections were stained for CK19, PDPN and αSMA. Representative images are displayed. Scale bar: 125 µm. **i**, Quantification of positive cells in (**h**), expressed as number of cells per mm² of tumor (control n=6, *Serpinb2* KO n=6, *Serpine1* KO n=6). **j**, Representative H&E and Masson’s trichrome collagen stainings (in blue) of representative tumor sections from orthotopically transplanted control, *Serpinb2* KO, and *Serpine1* KO mice. Scale bar: 50 µm. **k**, Quantification of trichrome-stained collagen (**j**) as a percentage of the tumor tissue area (control n=4, *Serpinb2* KO n=4, *Serpine1* KO n=4). Statistical significance in **a**, **c**, **e**, **f**, **g**, **i** and **k** was determined using a two-tailed, unpaired Student’s t-test. The p value for **d** was calculated using the log-rank (Mantel-Cox) test. TME: tumor microenvironment, KO: knock-out.

To better understand the functions of *Serpinb2* and *Serpine1* in PDAC progression, we generated individual KOs of each gene in KPC cells using two distinct sgRNAs, then confirmed successful gene deletion by western blot analysis (**Figure S6a**). Neither gene KO had an impact on KPC cell proliferation *in vitro* (**Figure S6b**), consistent with our pooled *in vitro* analysis and publicly available DepMap data (**Figure S3b**, **S3c**).

We orthotopically transplanted *Serpinb2*, *Serpine1*, or control KO KPC cells into syngeneic, immunocompetent mice (**Figure 3b**). Consistent with our Perturb-map results, individual KOs of *Serpinb2* and *Serpine1* significantly slowed tumor growth *in vivo* compared to control KO, reducing mean tumor burden by more than 50% at comparable time points (**Figures 3c** and **S6c**). This translated to an extension of animal survival, with some mice living more than 10 days longer than control, which is substantial considering the aggressiveness of the model (**Figures 3d**). These findings confirm an important functional role for both Serpins in promoting tumor growth *in vivo*.

Perturb-map analysis had found an increase in CD8+ T cells proximal to *Serpinb2* and *Serpine1* KO clones. To validate these findings, we collected tumors from a cohort of mice, and analyzed immune cell infiltrates by flow cytometry. Both *Serpinb2* KO and *Serpine1* KO tumors had reduced numbers of macrophages (**Figure 3e**), with *Serpinb2* KO tumors also showing a decrease in neutrophils (**Figure 3f**). Analysis of the T cell compartment revealed a doubling of CD8+ T cells in the Serpin KO tumors (**Figure 3g**), consistent with the enrichment we detected in the heterogeneous Perturb-map context. There was also a trend toward decreased CD4+ T cells in KO tumors (**Figure S6d**).

To determine if stromal cell composition was also affected by loss of either Serpin, we stained tumors sections for specific cancer associated fibroblast (CAF) markers. Imaging revealed a reduction in the overall number of CAFs, identified as PDPN positive cells, and myofibroblastic CAFs (myoCAFs), defined by dual PDPN+ and αSMA+ staining, in *Serpine1* KO PDACs (**Figures 3h, 3i**). This reduction coincided with decreased collagen deposition, as determined by Masson’s trichrome staining (**Figures 3j**, **3k**). Interestingly, *Serpinb2* KO tumors also exhibited reduced collagen deposition, consistent with the *Serpine1* KO phenotype, potentially reflecting their shared roles in regulating plasminogen activation, fibrinolysis and MMP activation, processes known to influence collagen deposition and ECM remodeling^3,23–26^.

### *Serpinb2* and *Serpine1* promote PDAC resistance to anti-PD1 immunotherapy

To investigate how PDAC-derived PAI1 and PAI2 influence the transcriptional state of the TME within pancreatic tumors, we performed scRNA-seq on cells isolated from *Serpinb2*, *Serpine1* and control KOs. To define cell populations, we combined the data from both KOs and control tumors, representing a total of 15,233 cells analyzed. In all conditions we identified tumor cells, CAFs, acinar cells, endothelial cells, myeloid cells including macrophages, neutrophils and dendritic cells, B cells, plasma cells and T and NK cells (**Figure S7a**).

Clustering and annotation of T cell populations revealed a 3.5-fold and 1.7-fold reduction in terminally exhausted T cells in *Serpinb2* and *Serpine1* KO tumors, respectively, and a concomitant increase in CD8 effector T cells (**Figure 4a-c**, **S7a-c**). To validate these findings, we stained tumor tissue sections for CD8 and Granzyme B (GZMB), a key mediator of cytotoxic T cell activity. The loss of either *Serpinb2* or *Serpine1* resulted in a significant increase in the number of CD8+ T cells within the tumors, including a notable increase in GZMB+ CD8+ T cells (**Figure 4d**, **4e**). These results indicate that *Serpinb2* and *Serpine1* activity promotes T cell exhaustion and/or effector exclusion, and further suggested that they promoted tumor growth by suppressing the adaptive immune response. Therefore, targeting *Serpinb2* and *Serpine1* may reinvigorate anti-tumor immunity in PDAC. To test this, we transplanted *Serpinb2*, *Serpine1*, or control KO KPC cells into Rag2-/- mice. As observed with Perturb-map, KO of these regulators of fibrinolysis resulted in faster tumor growth (**Figure 4f-g**). This confirms that in the context of immune pressure, cancer-derived PAI1 and PAI2 are acting to suppress anti-tumor immunity.

**Figure 4.**
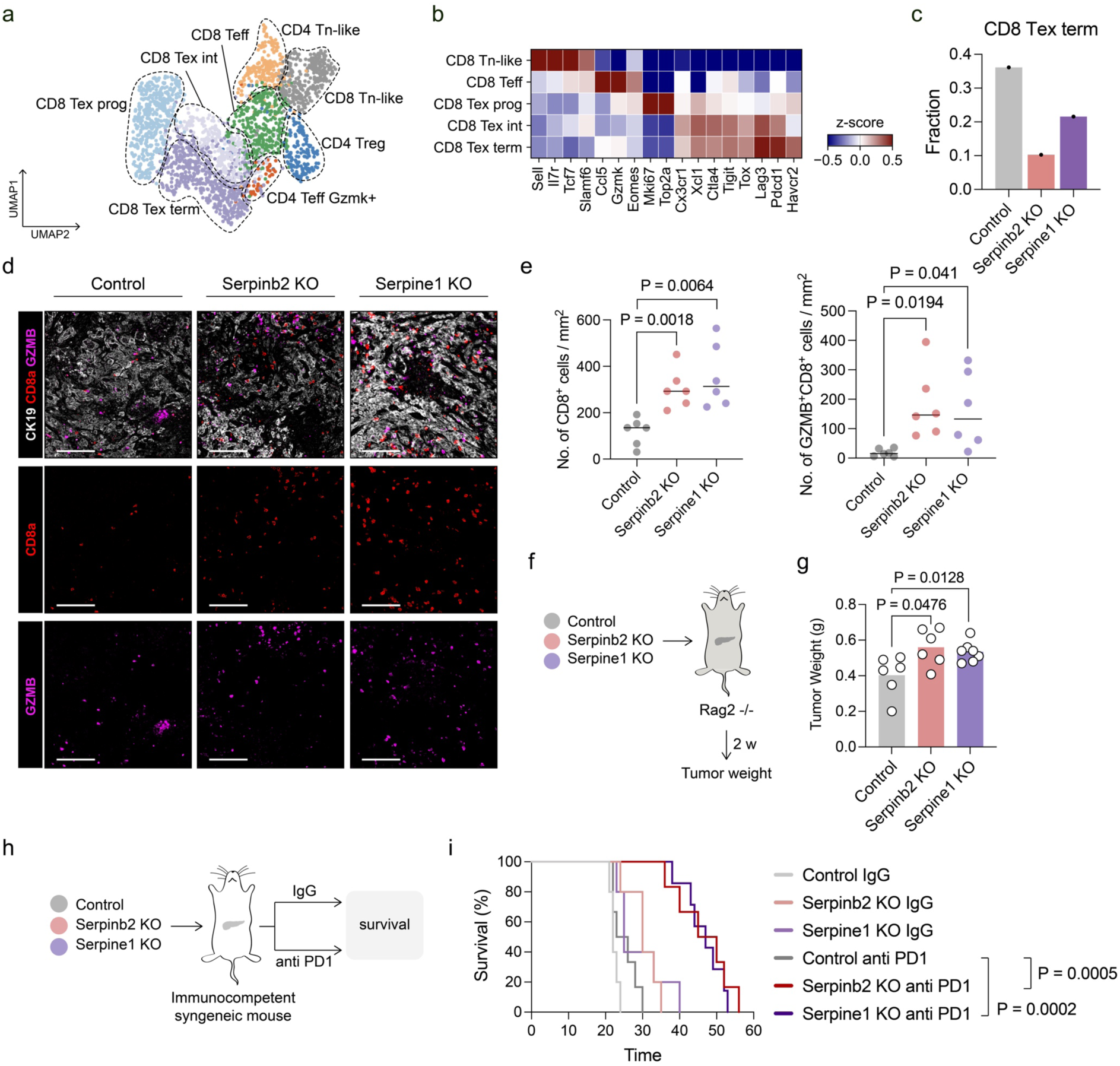
Knock-out of *Serpinb2* and *Serpine1* triggers T cell-mediated anti-tumor immune responses. **a**, UMAP projection of T cells with labeled subsets across control, *Serpinb2* KO, *Serpine1* KO. **b**, Heatmap showing z-score of hallmark genes across the identified CD8 T cell subsets. **c**, Relative frequency of CD8 terminally exhausted subsets across the investigated conditions. **d**, Pseudocolored immunohistochemistry of tumors from control, *Serpinb2*, and *Serpine1* KOs, isolated two weeks post-transplantation. Sections were stained for CK19, CD8a and GZMB. Representative images are displayed. Scale bar: 125 µm. **e**, Quantification of positive cells in (**d**), expressed as number of cells per mm² of tumor (control n=6, *Serpinb2* KO n=6, *Serpine1* KO n=6). **f**, Control, *Serpinb2* KO, and *Serpine1* KO tumor cells were orthotopically transplanted into immunodeficient Rag2-/- mice. Phenotypic analysis was performed on tumors two weeks post-transplantation. **g**, Tumor weight comparison between control, *Serpinb2* KO, and *Serpine1* KO tumors isolated from Rag2-/- mice (control n=6, *Serpinb2* KO n=6, *Serpine1* KO n=7). **h**, Scheme of the experimental set-up used to investigate the synergy between *Serpinb2*, *Serpine1* deletion and anti-PD1 treatment. **i**, Kaplan–Meier survival curves of mice with orthotopic control, *Serpinb2*, and *Serpine1* KOs tumors, IgG or anti-PD1 treated. 5-7 mice per condition were analyzed. Statistical significance in **e** and **g** was determined using a two-tailed, unpaired Student’s t-test. The p value for **i** was calculated using the log-rank (Mantel-Cox) test. KO: knock-out, scRNA-seq: single cell RNA sequencing.

To determine if *Serpinb2* and *Serpine1* are also functioning in PDAC resistance to immunotherapy, we transplanted KO KPC cells into the pancreas of immunocompetent mice, and starting 10 days post-transplantation, animals were treated with either an IgG control or anti-PD1 antibody (**Figure 4h**). Consistent with clinical observations in PDAC patients, anti-PD1 monotherapy had little to no impact on the growth of wildtype tumors. Strikingly, in tumors lacking *Serpinb2* or *Serpine1*, anti-PD1 treatment resulted in a major reduction in tumor growth, as evident by the significant increase in survival. Deletion of either gene combined with anti-PD1 therapy nearly doubled the median survival time, from 24.5 days in control KPC tumors treated with anti-PD1, to 47.5 days and 47 days in *Serpinb2* KO and *Serpine1* KO tumors, respectively (**Figure 4i**). These findings demonstrate that *Serpinb2* and *Serpine1* deletion acts synergistically with anti-PD1 therapy, highlighting a potential therapeutic strategy for overcoming resistance to immune checkpoint inhibitors in PDAC.

### *Serpinb2* and *Serpine1* deletion alters PDAC cells intrinsic immune and stromal regulatory programs

To investigate the mechanisms driving changes in the TME, we performed transcriptomic analysis of control, *Serpinb2* KO, and *Serpine1* KO tumor cell lines (**Figure S8a**). Though KO of the Serpins did not affect tumor growth *in vitro*, it led to distinct transcriptional changes in the cells (**Figure S8b**). The major role ascribed to both Serpins is extracellular, however intrinsic cell effects have also been described for *Serpine1*^27^, which may relate to an intracellular function of these proteins or reflect alterations in extracellular conditions mediated by loss of PAI1 or PAI2 release, as urokinase and other substrates of these inhibitors are produced by cells in culture. Considering the observed phenotypic changes in the TME, we focused on genes with extracellular functions and identified both shared and unique changes between the two KOs (**Figure S8c**). Among the shared downregulated genes was *Thbs1*, a matricellular protein implicated in fibrinolysis and TME modulation via activation of latent TGF-β within the extracellular space^28^, suggesting that both Serpins influence extracellular matrix dynamics. Unique to *Serpinb2* KO cells was the downregulation of *Cxcl3* and *Cxcl5*, chemokines associated with neutrophil recruitment. In contrast, *Serpine1* KO tumor cells exhibited reduced cancer cell-derived expression of *Tgfb1*, a key regulator of immunosuppression and modulator of CAF states^29^ (**Figure S8d**). This finding aligns with the observed reduction in CAF abundance and state differences in *Serpine1* KO tumors and underscores the role of *Serpine1* in promoting a CAF-rich, immunosuppressive TME. Together, these results suggest that *Serpinb2* and *Serpine1* play a role in influencing distinct transcriptional programs of extracellular factors produced by PDAC cells, and contribute to the immune and stromal composition of the PDAC microenvironment.

### Tumor macrophages accumulate in fibrin(ogen)-rich regions and play a central role in mediating SERPINB2- and SERPINE1-driven immunosuppression

The coagulome, which encompasses the interplay between coagulation and fibrinolysis pathways, has been increasingly recognized for its role in shaping immune suppression and stromal remodeling within the TME^30^. However, despite its acknowledged importance, the precise mechanisms underlying these effects remain poorly defined, particularly concerning the role of plasminogen activation inhibitors like *SERPINB2* and *SERPINE1*. These inhibitors physiologically block the conversion of plasminogen to plasmin, thereby preventing fibrin clot degradation. Given *SERPINB2* and *SERPINE1* increased expression in PDAC, we examined fibrin(ogen) deposition in the ECM and the effects of *Serpinb2* and *Serpine1* KO on ECM composition.

In orthotopic PDAC mouse tumor sections, we identified fibrin(ogen)-dense areas (**Figure 5a-c**), as previously reported^31^. Comparison between control and Serpin KO tumors revealed a significant reduction in fibrin(ogen)-stained regions, indicating that both *Serpine1* and *Serpinb2* promote fibrin deposition in PDAC (**Figure 5a-c**). As macrophages and neutrophils directly bind fibrin in inflammatory contexts via the integrin αMβ2 (composed of Cd11b and Cd18), we sought to determine if the reduction of these cells in *Serpine1* and *Serpinb2* KO tumors might be related to fibrin(ogen) deposition. We stained both mouse and human tissues for fibrin(ogen) as well as the two Serpins, and markers of cancer cells and defined immune cell types (**Figure 5c**). To minimize border effects of immune cells, we restricted our analysis to central tumor areas (**Figure S9a**). Interestingly, within fibrin(ogen)-dense areas there was a reduction in the distance between tumor macrophages and CK19+PAI1+ and CK19+PAI2+ tumor cells (**Figure 5c-e**). Neutrophils were also more proximal to CK19+PAI2+ cells, though not CK19+PAI1+ tumor cells (**Figure S9b**), in line with the observed selective transcriptional regulation of neutrophil chemoattractant expression in PDAC cells by PAI2 (**Figure S8**). These spatial associations were corroborated in human PDAC samples, highlighting the conserved nature of these interactions (**Figure S9c-e**). These findings indicate that macrophages and neutrophils, both of which express the fibrin(ogen)-binding integrin αMβ2, preferentially localize within fibrin(ogen)-rich areas of tumors, and suggest that this is mediated by cancer cell production of *Serpine1* and *Serpinb2*.

**Figure 5.**
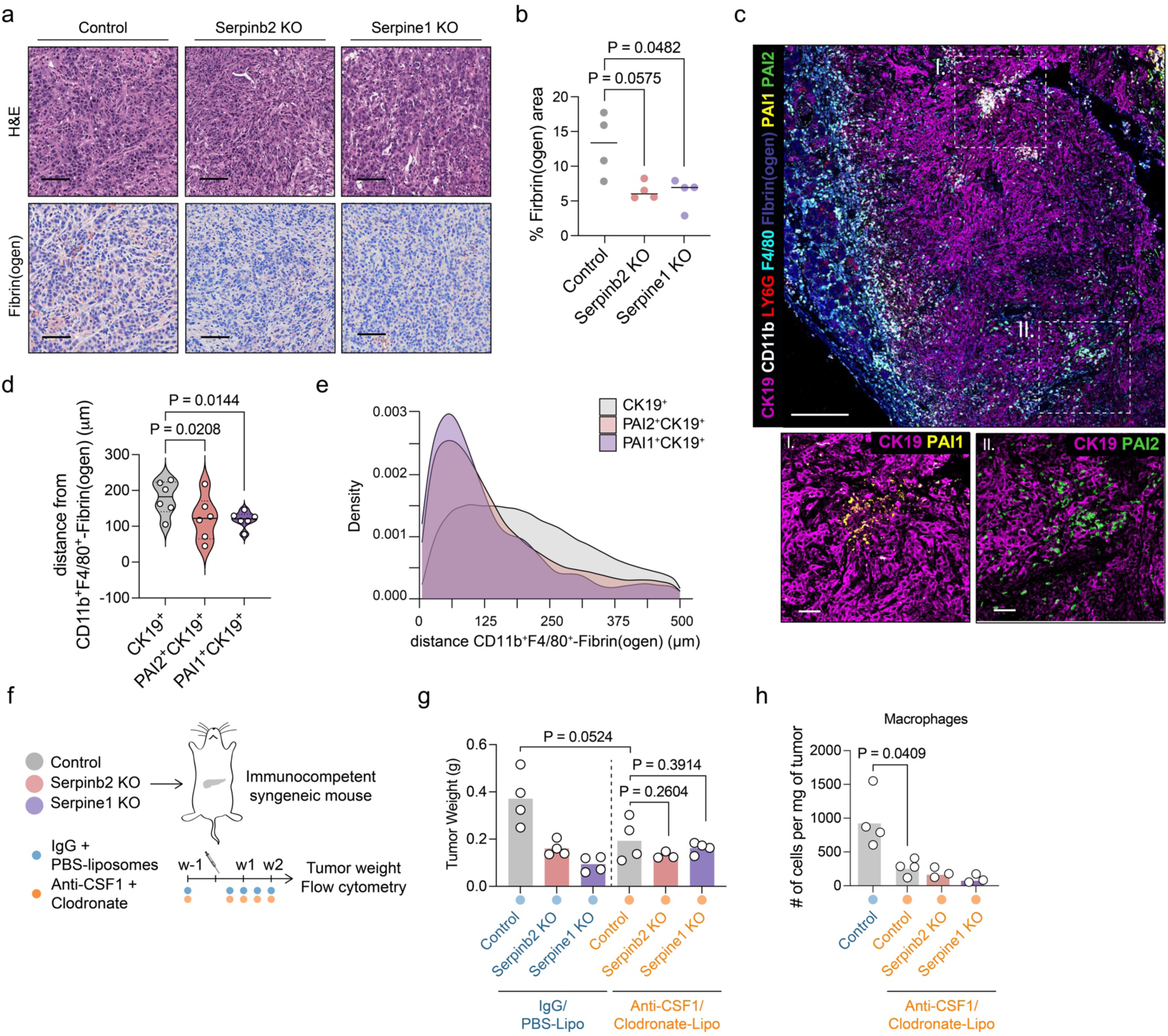
Tumor associated macrophages localize in fibrin(ogen) dense areas. **a**, Representative H&E and Fibrin(ogen) IHC staining of tumor sections from orthotopically transplanted control, *Serpinb2* KO, and *Serpine1* KO mice. Scale bar: 50 µm. **b**, Quantification of Fibrin(ogen) IHC as a percentage of tissue area (control n=4, *Serpinb2* KO n=4, *Serpine1* KO n=4). **c**, Pseudocolored immunohistochemistry of tumors from control mice, isolated two weeks post-transplantation. Sections were stained for CK19, CD11b, LY6G, F4/80, Fibrin(ogen), PAI1, and PAI2. Representative images are shown. Scale bar: 200 µm for the main image and 50 µm for insets. **d**, Comparison of distances between CD11b+F4/80+ cells co-localizing with Fibrin(ogen) (CD11b+F4/80+/Fibrin(ogen)), and CK19+, PAI1+CK19+, and PAI2+CK19+ cells in control tumors (n=6). Only the tumor center area, as visualized in figure **S9**, panel (**a**), was used for the analysis. **e**, Density plot showing the spatial distribution of CK19+ (grey), PAI2+CK19+ (red) and PAI1+CK19+ (purple) cells relative to CD11b+F4/80+/Fibrin(ogen) immune cells. The x-axis shows the distance (μm) from CD11b+F4/80+/Fibrin(ogen) cells and the y-axis the distribution density. Only the tumor center area, as visualized in figure **S9**, panel (**a**), was used for the analysis. **f**, Schematic of the experimental design. Mice were pre-treated with IgG+PBS-liposomes or anti CSF1+clodronate and orthotopically transplanted with control, *Serpinb2* KO, or *Serpine1* KO KPC cells. Mice were treated twice weekly for two weeks. Tumors were harvested for phenotypic analysis. **g**, Tumor weight comparison between IgG+PBS-liposome control (n=4), *Serpinb2* KO (n=4), and *Serpine1* KO (n=4) tumors, and anti CSF1+clodronate-liposome control (n=4), *Serpinb2* KO (n=3), and *Serpine1* KO (n=4) tumors. **h**, Flow cytometry analysis of tumors isolated from IgG+PBS-liposome control, and anti CSF1+clodronate-liposome control, *Serpinb2*, and *Serpine1* KO mice two weeks after orthotopic transplantation. The absolute numbers of macrophages are shown (IgG+PBS-liposome control n=4, anti CSF1+clodronate-liposome control n=4, anti CSF1+clodronate-liposome *Serpinb2* KO n=3, anti CSF1+clodronate-liposome *Serpine1* KO n=3). Statistical significance in **b**, **g** and **h** was determined using a two-tailed, unpaired Student’s t-test. P value in **d** was calculated with a two-tailed, paired t-test KO: knock-out, IHC: immunohistochemistry

To determine if the Serpins were protecting the tumors through an effect on macrophages, we implanted mice with control or Serpin KO tumors and treated them with anti-CSF1 and clodronate to deplete macrophages (**Figure 5f**). In mice treated with isotype control antibody and PBS-liposomes, we once again found that *Serpinb2* and *Serpine1* KO tumor growth was significantly reduced (**Figure 5g**). However, when macrophages were depleted (**Figure 5h**), there was no difference in tumor burden between the control and KO tumors (**Figure 5g**), indicating no additive effects when macrophage depletion was combined with *Serpinb2* or *Serpine1* KO. This suggests that macrophages are the primary mediators of *Serpinb2*-and *Serpine1*-induced immunosuppression.

### PDAC-derived PAI1 and PAI2 promotes the enrichment of M2-like Marco macrophages that exhibit a fibrin(ogen) binding gene signature

ScRNA-seq revealed extensive transcriptional reprogramming of the macrophages in *Serpinb2* and *Serpine1* KO tumors (**Figure 6a-c** and **Figure S10a, S10b**), highlighted by the enrichment of gene expression signatures associated with inflammatory responses, interferon signaling, and cytokine production pathways (**Figure 6b**). Notably, *Serpinb2* and *Serpine1* KOs induced largely concordant changes in macrophages, with an increased abundance of Cxcl9+ populations. We observed also a reduction in Marco+ macrophages in *Serpine1* KO tumors, and fewer proliferating macrophages in *Serpinb2* KOs (**Figure 6c**). These findings suggest a remodeling of the tumor microenvironment toward an immune-activated state, consistent with studies linking Cxcl9+ macrophages to T cell recruitment, favorable prognosis and immunotherapy response^32,33^, and Marco+ macrophages to immune suppression^13,34^.

**Figure 6.**
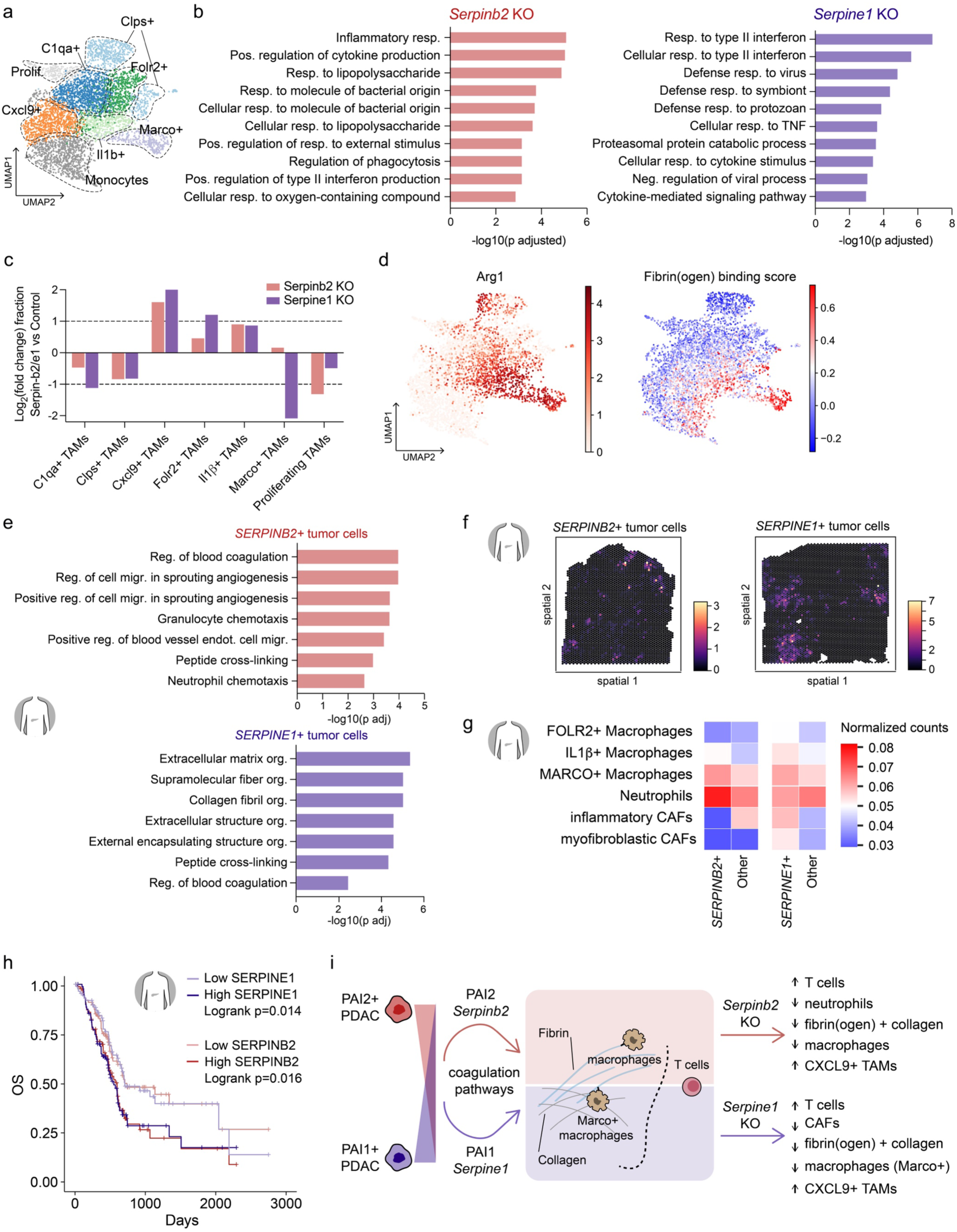
*Serpinb2* and *Serpine1* expression modulates macrophage subsets and correlates with distinct TME landscapes and poor overall survival in hPDAC. **a**, Control, *Serpinb2* KO, and *Serpine1* KO tumors were subjected to scRNA-seq analysis. UMAP projection of macrophages with annotated subsets across control, *Serpinb2* KO, *Serpine1* KO tumors. **b**, Displayed are the top 10 gene ontology (GO) “biological process” gene sets enriched in macrophages infiltrating *Serpinb2* and *Serpine1* KO tumors. **c**, Relative abundance of macrophage subsets in the two KO conditions compared to control. The log2 fold change in abundance relative to control for all populations is shown. **d**, UMAP projection of *Arg1* expression in macrophages (left) and of “fibrin(ogen) binding score” (right). **e**, Tumor cells, from the hPDAC dataset Cui Zhou, et al.^34^, expressing *SERPINB2* or *SERPINE1* were subjected to gene set enrichment analysis. The top 7 GO “biological process” gene sets enriched in these tumor cells are shown. **f**, Proportions of macrophages, neutrophils, and CAFs in the local TME of *SERPINB2*+ and *SERPINE1*+ tumor Visium ST spots. Visium ST spots were deconvoluted using cell2location using the Cui Zhou, et al.^34^ scRNA-seq reference dataset. Spots were classified as positive for *SERPINB2* or *SERPINE1* if they contained at least four tumor cells and met minimum gene expression thresholds (see methods for details). **g**, Analysis of cell type infiltration in *SERPINB2*+ (left) and *SERPINE1*+ (right) tumor regions. Tumor spots identified via Visium ST were stratified based on *SERPINB2* and *SERPINE1* expression. The number of immune and stromal identified cell populations in the local TME of *SERPINE1*+ and *SERPINB2*+ tumor cells were compared to the TME of tumor cell spots lacking their expression. **h**, Kaplan–Meier survival analysis comparing the overall survival of patients with either low or high tumor expression of *SERPINB2* and *SERPINE1*. **i**, Scheme of the proposed role for *SERPINB2* or *SERPINE1* function in PDAC. The p value in **h** was calculated using the log-rank (Mantel–Cox) test. TME: tumor microenvironment, hPDAC: human PDAC, scRNA-seq: single cell RNA sequencing, ST: spatial transcriptomics.

To further explore the functional impact of fibrin(ogen) on macrophage phenotype, we calculated a fibrin(ogen) binding score using genes from the “fibrinogen binding” gene ontology molecular function and projected it on the macrophage cluster. Quite strikingly, the *Arg1* expressing macrophages, which included the Marco+ cluster, specifically upregulated the “fibrinogen binding” gene signature (**Figure 6d**). This, together with the imaging data, suggests that immunosuppressive macrophages localize in fibrin dense areas, and that *Serpinb2*- and *Serpine1*-driven fibrin-rich ECM shapes the immunosuppressive TME. This reveals that the PAI1/PAI2/fibrin(ogen) axis plays a previously unrecognized role in recruiting and polarizing tumor-associated macrophages, fostering immune dysfunction in PDAC.

### *SERPINB2* and *SERPINE1* expression in human PDAC is linked to distinct subtypes, TME landscapes and poor overall survival

To place our findings into the heterogenous TME landscape of human PDAC, we next analyzed *SERPINB2* and *SERPINE1* expression across a collection of human PDAC scRNA-seq and matched Visium spatial transcriptomics (ST) datasets^35^. There, we identified subsets of cancer cells that expressed either *SERPINB2* or *SERPINE1* (**Figure S11a**), each associated with common and distinct transcriptional profiles. Both groups of tumor cells expressed gene signatures related to negative regulation of fibrinolysis and positive regulation of blood coagulation (**Supplementary table 1**). *SERPINB2*-high cells were enriched for gene ontology (GO) terms related to granulocyte and neutrophil chemotaxis (**Figure 6e**), consistent with the inflammatory phenotypes observed in our orthotopic models. In contrast, *SERPINE1*-high tumor cells displayed gene signatures linked to ECM remodeling, and TGF-β signaling (**Figure 6e**, **Supplementary table 1**), corroborating our mouse *in vivo* data implicating CAFs and collagen deposition. Spatial deconvolution of Visium ST data revealed that immunosuppressive macrophages (IL1β+ and MARCO+) were preferentially localized near tumor cells expressing either *SERPINB2* or *SERPINE1*, with neutrophils enriched in proximity to *SERPINB2*-expressing cells and CAFs predominantly associated with *SERPINE1*-expressing cells (**Figure 6f**, **6g**).

Considering the heterogeneity of *SERPINB2* and *SERPINE1* expression, we next mapped PDAC transcriptional subtypes^36,37^ onto our analyzed dataset and identified distinct associations for *SERPINE1* and *SERPINB2* expression. Tumor cells with high *SERPINE1* expression exhibited preferential features of the basal-like PDAC subtype, whereas *SERPINB2*-high tumor cells were characterized by classical transcriptional signatures (**Figure S11b, S11c**).

Because both genes promoted immunosuppression and pancreatic tumor growth in our models, we hypothesized that their expression might predict patient outcomes. Despite the fact that both genes were expressed by a relatively small fraction of cancer cells (in both mouse and human PDAC tumors), survival analysis based on bulk RNA-seq data indicated that PDAC patients with low expression of both *SERPINB2* and *SERPINE1* had significantly better overall survival (**Figure 6h**). This trend was further validated using the TCGA pan-cancer dataset, where low expression of these genes correlated with reduced mortality risk in several cancers, including pancreatic cancer (**Figure S11d**).

Considering the limited availability of clinical trial data evaluating immune checkpoint blockade in PDAC, and our findings highlighting the relevance of *SERPINB2* and *SERPINE1* in other cancer types, we analyzed data from the JAVELIN Renal 101 trial, which tested the combination of avelumab and axitinib versus sunitinib in advanced renal cell carcinoma^38^. Within the avelumab plus axitinib cohort, we stratified patients based on high versus low expression of *SERPINB2* and *SERPINE1*. Patients with high *SERPINE1* expression exhibited significantly poorer responses to the combination treatment with ICB (**Figure S11e**). For *SERPINB2*, a slight trend was observed, though it did not reach statistical significance (**Figure S11e**).

Collectively, these findings support a model in which cancer cell-derived PAI2 (*SERPINB2*) and PAI1 (*SERPINE1*) drive local ECM and tumor immune landscape remodeling, facilitating the recruitment of pro-tumorigenic and immunosuppressive cells, and CD8+ T cell exclusion and dysfunction thereby promoting PDAC progression and more aggressive tumor phenotypes (**Figure 6i**).

## Discussion

Here, we conducted a CRISPR screen targeting a focused set of receptors and ligands expressed by PDAC cells, predicted to influence the extracellular milieu and immunosuppressive properties of the tumor. A key challenge in such screens is compensatory effects from neighboring cells lacking the same genetic perturbation. To address this, we utilized Perturb-map, a platform that enables spatial resolution of CRISPR-edited cells in tissue through multiplex imaging^12,13^. This system enabled us to observe spatial and compositional changes in heterogeneous clonal growth and the TME over time, which has not previously been done. Our initial gene set comprised candidates expressed by tumor cells and implicated in promoting TME organization and immune suppression functions. Many of these genes were indeed critical *in vivo*, but not *in vitro*, as their knock-out resulted in reduced or increased abundance relative to internal controls by tumor endpoint. However, our data also highlight the remarkable plasticity of tumor cells, which adapt to their environment and reprogram the extracellular milieu over time, enabling in some instances continued clonal expansion (e.g., *Fst*, *Serpinf2*, *Lta4h*) only after a certain period of time. In contrast, others, such as knock-outs of *Serpinb2* and *Serpine1*, showed strong and consistent fitness defects across time points along with a profound remodeling of the TME, which were apparent even within a highly heterogeneous milieu of different gene KO clones.

Our use of Perturb-map provided unique insights into the spatial dynamics of clonal evolution and the localized remodeling of the TME. The multiplex imaging approach, combined with neighborhood enrichment analyses, revealed gene-specific effects on immune cell distribution within the tumor mass. For example, *Serpinb2* and *Serpine1* knock-outs were associated with increased CD8+ T cell density and reduced macrophages, indicative of a less immunosuppressive microenvironment. Temporal analyses showed that neighborhood patterns were largely established by day 14, suggesting that early TME remodeling events likely influenced later-stage clonal dominance. Strikingly, despite the inherent heterogeneity of the PDAC TME, Perturb-map revealed consistent and reproducible immune cell distribution patterns around specific gene KO clones, underscoring the robustness of gene-specific effects in shaping the immune landscape. These findings demonstrate how individual genes regulate clonal fitness by influencing the surrounding immune environment, with important implications for therapeutic targeting of the tumor-host interface.

A notable clinical feature of PDAC is the pervasive activation of the coagulation system, which contributes to tumor progression and significantly impacts patient morbidity^39^. Among all malignancies, PDAC exhibits one of the highest incidences of cancer-associated venous thromboembolism^40^. Within the TME, procoagulant conditions exacerbate hypoxia and nutrient deprivation while reshaping the ECM through fibrin deposition and crosslinking, leading to marked alterations in the tissue’s biophysical properties^30^. Counterbalancing the coagulation cascade is the fibrinolytic system, whose dysregulation has been linked to chronic inflammatory diseases, such as neurological and mucosal inflammations^41,42^. At the core of this system, plasmin serves as a key enzyme in fibrinolysis, facilitating the clearance of fibrin-rich matrices in both vascular and extravascular compartments.

*Serpine1* (PAI1) and *Serpinb2* (PAI2), are key players in the fibrinolytic pathway, as they inhibit plasmin activators and therefore fibrinolysis. Our analysis of *Serpine1* and *Serpinb2* function identified them as important regulators of several key components of the PDAC ecosystem. Previous reports have hinted at *Serpine1* function in shaping the PDAC TME^43^, however the mechanistic contribution of both genes was so far unknown. We find that *Serpinb2* and *Serpine1* loss impaired tumor growth by the depletion and reprogramming of immunosuppressive macrophages and the enrichment of cytotoxic CD8+ T cells. Mechanistically, this appeared linked to a PAI1 and PAI2 instructed fibrin- and collagen-rich ECM that recruits macrophages via fibrin-binding receptors like αMβ2 integrin (CD11b/CD18), fostering local immunosuppression, as observed for neutrophils in oral mucosal barrier dysfunctions^42^. Fibrinolysis is closely tied to the infiltration and activity of tumor associated macrophages across multiple cancer types, a connection that was suggested to help explain the prognostic relevance of fibrinolysis proteins, including PAI1^44,45^. In large transcriptomic analyses, *SERPINE1* mRNA levels consistently correlate with the macrophage marker CD163, further underscoring this link^44^. These data support our findings showing that dysregulated fibrinolysis contributes to immunosuppression and stromal reprogramming through macrophage-mediated mechanisms.

While the role of the ECM as a barrier to immunotherapy in PDAC is well-recognized^46^, our findings provide novel insights into how specific ECM components, directed by *Serpinb2* and *Serpine1*, actively shape the immunosuppressive microenvironment. Collagens can provide physical restriction to tumor entry, and loss of the Serpins led to reduce collagen density. However, the enrichment of macrophages in fibrin(ogen) depots and expression of the fibrin-induced signature in *Arg1*-expressing macrophage, strongly suggests that *Serpinb2* and *Serpine1* control of fibrin(ogen) deposition in the ECM also had a direct effect on polarizing and retaining immunosuppressive macrophages around cancer cells.

Pancreatic tumors lacking *Serpinb2* and *Serpine1* exhibited a synergistic response to PD1 checkpoint blockade, whereas this therapy was ineffective in control tumors with intact ECM regulation. This identifies two specific mediators of ECM remodeling that promote immunotherapy resistance. TGF-β is another factor established to be a key regulator of the TME, ECM deposition, and PD1 resistance^47^. Notably, in pancreatic cancer cells lacking either *Serpinb2* or *Serpine1* there was downregulation of *Thbs1*, a pivotal activator of latent extracellular TGF-β1^28^. This suggests that *Serpine1* and *Serpinb2* may not only suppress anti-tumor immunity by altering fibrin homeostasis, but also via activation of TGF-β signaling. Interestingly, *Serpine1* is a transcriptional target of the TGF-β pathway^47^, pointing to a positive feedback loop between *Serpine1* and *Serpinb2* expression, TGF-β signaling, and CAF activation that orchestrates the TME towards tumor promotion and immunosuppression. TGF-β induced *Serpine1* expression in turn prevents degradation of ECM components by extracellular proteases leading to a collagen-rich stroma that impairs T cell infiltration^3,47^. The coordinated decrease of *Thbs1* and *Tgfb1* upon *Serpinb2* and *Serpine1* loss suggests that these genes function to integrate ECM remodeling with TGF-β-driven programs, enabling immune evasion and stromal activation. These findings position the fibrinolytic system as a key mediator of TME dynamics, linking structural ECM cues to immunosuppressive signaling.

The expression of *Serpinb2* and *Serpine1* was heterogeneous across PDAC patients, with a notable association between *Serpinb2* and the classical subtype, and *Serpine1* with the basal-like subtype. This, along with the fact that the two Serpins were generally not detected in the same cancer cells, implies context-specific activity of *Serpine1* and *Serpinb2* and provides insights into their role in shaping the distinct TME features of the two PDAC subtypes. Both Serpins regulated distinct gene expression programs within the cancer cells themselves, which may relate to the subtype associations. Though neither affected cancer cell fitness *in vitro*, they both regulated TME and ECM composition and tumor progression *in vivo*, which differed between immunocompromised and immunocompetent contexts.

These studies point to a dual mechanism by which *Serpinb2* and *Serpine1* shape the tumor ecosystem: through direct modulation of fibrinolysis and through intracellular transcriptional rewiring that drive immune and stromal dynamics. This dual role positions these genes as central mediators of TME remodeling, instructing key drivers of tumor progression, TAMs, neutrophils, CAFs, the ECM and T cell dysfunction, with implications for a dual therapeutic strategy aimed at targeting fibrinolytic pathways and reprogramming the TME globally to support anti-tumor immunity. Together with our data demonstrating poor prognosis and immunotherapy resistance of human PAI1 (*SERPINE1*) and PAI2 (*SERPINB2*) rich tumors, this indicates that both Serpins are prime targets for future therapeutic interventions.

## Acknowledgments

We thank members of the Brown lab for thoughtful feedback on the manuscript, members of the Center for Comparative Medicine and Surgery at the Icahn School of Medicine at Mount Sinai for animal husbandry and members of the Human Immune Monitoring Center at the Icahn School of Medicine at Mount Sinai for scRNA-seq assistance. C.F. was supported by the Cancer Research Institute Irvington Postdoctoral Research Fellowship (award no. CRI4641). A.T. was supported by NIH F30CA287690. B.D.B. and M.M. were supported by U01CA282114, R01CA254104, R01CA257195 and funding from the Feldman Foundation. Computational resources at the Icahn School of Medicine supported by NIH UL1TR004419.

## Author contributions

C.F., S.R.N. and B.D.B. conceptualized the project. C.F., S.R.N. and B.D.B. designed experiments. C.F., S.R.N., A.T., D.C., H.T.P, M.D., G.M. and M.D.P performed experiments. C.F., M.M.S., B.S. and M.D.P. performed data analysis. M.M., A.B. and B.D.B. provided intellectual input, essential reagents and computational tools. C.F. wrote the manuscript. B.D.B. edited the manuscript. All authors provided feedback on the manuscript draft.

## Methods

### Lentiviral vector construction and production

A comprehensive protocol for generating Pro-Code/CRISPR lentiviral vectors and library pools is available on the Addgene website, as part of the Pro-Code vector kit (Addgene #1000000197) and has been described previously. Below, we provide a brief overview of the procedure. For selecting sgRNA sequences targeting each gene, we used the Brie CRISPR library for mouse genes^48^. A complete list of the oligonucleotides used for the experiments in this paper is provided in **supplementary table 2**. Library cloning was performed using nuclear Pro-Code/CRISPR lentiviral vectors (Addgene #1000000197), while validations utilized lentiCRISPRv2 (Addgene #52961, ^49^). Library cloning was conducted in a 96-well plate format, where constructs were annealed, ligated, and transformed individually but processed in parallel, as outlined in the Pro-Code vector kit protocol. Oligonucleotides were prepared by resuspending them to 100 µM in water. For annealing, forward and reverse oligos were mixed to a final concentration of 2 µM, combined with 10X NEBuffer 2.1 and water. Next, they were heated to 95°C for 5 minutes, followed by gradual cooling to room temperature. The Pro-Code/CRISPR lentiviral vector was digested with BbsI-HF (NEB), as per the manufacturer’s instructions, and purified using Qiagen PCR purification columns. Annealed oligos were ligated into the digested vector backbone by combining 50 ng of digested plasmid with 6–8 ng of annealed oligos, incubating at room temperature for 10 minutes with Quick Ligase (NEB). Next, 5 µL of the ligation reaction was transformed into 50 µL of TOP10 chemically competent bacteria. After incubating on ice for 30 minutes, the bacteria were heat-shocked at 42°C for 30 seconds, cooled on ice for 2 minutes, and plated on LB ampicillin agar plates overnight at 37°C. Colonies were picked, cultured, and plasmid DNA was extracted using the Zymo ZR™ Plasmid MiniPrep Classic Kit. The sgRNA sequence was confirmed via Sanger sequencing. For cloning into the lentiCRISPRv2 backbone, a similar process was followed. However, sgRNAs were inserted into the BsmBI site.

Lentiviral vector production was performed as previously described^50^ and detailed in the Pro-Code_Kit_Methodology.pdf on the Addgene website. Briefly, 293T cells were seeded at 500,000 cells per well in 6-well plates and incubated at 37°C with 5% CO2. After 24 hours, cells were transfected using calcium phosphate with third-generation lentiviral packaging plasmids and the transfer plasmid [pVSV (1 µg), pMDLg/pRRE (2 µg), pRSV-REV (1 µg), and Pro-Code/CRISPR vector (6 µg)]. The plasmids were first mixed with 2.5 M CaCl2, vortexed, and incubated for 10 minutes. Then, 2X HBS solution (281 mM NaCl, 100 mM HEPES, 1.5 mM Na2HPO4, pH 7.05) was added dropwise with gentle vortexing. The transfection mixture was applied to cells, and the media was replaced after 14 hours. Supernatants were harvested 30 hours after media replacement, filtered through a 0.22 µm PVDF disc filter, and stored at - 80°C.

### Cell culture

FC1245 PDAC cells (KPC cells, in this manuscript) were generated from a primary tumor in a *Kras^LSL-G12D/+;^ Trp53^LSL-R^*^172^*^H/+^; Pdx1-cre* mouse and were kindly provided by D. Tuveson (Cold Spring Harbor Laboratory).

These cells, and all lines resulting from their genetic modification, were routinely passaged in DMEM (Thermo Fisher Scientific) containing 10% heat-inactivated FBS and 100 U/ml penicillin/streptomycin, for no more than 25 to 30 passages. All cell lines were routinely tested for mycoplasma; PC+ cells were verified by flow cytometry (for mCherry positivity, as readout for PC presence in the culture) and mass cytometry (for their sgRNA/PC identities) before every *in vivo* experiment.

To determine doubling time of cell lines, 1000 cells per well were seeded in 100 μl of growth medium in technical triplicates in 96 well plates. After 24h intervals, cells were fixed and stained with 0.2% Crystal Violet in an ethanol:water solution. Crystal Violet was solubilized with 10% acetic acid and absorbance was quantified at 595 nm. The resulting values were used to determine doubling times. Experiments were performed in three biological replicates.

### Vector transduction

For the Perturb-map experiment, KPC cells were transduced with Pro-Code/CRISPR lentiviral vectors. Cells were seeded in 12-well plates at a density of 20,000 cells per well, 24 hours prior to transduction. The next day, lentiviral vectors were added to the cells in the presence of 5 µg/ml polybrene (Millipore) at a low multiplicity of infection (MOI), with each well receiving a distinct Pro-Code/CRISPR vector. The transduced cell populations were pooled based on their mCherry+ percentage to achieve an equal distribution of Pro-Code/CRISPR populations. Cells expressing PC/CRISPR were sorted for mCherry positivity, ensuring >99% purity. Subsequently, KPC cells were transduced with Cas9 lentivirus and selected using 4 µg/ml puromycin. PC/CRISPR cells, both with and without Cas9, were maintained in culture with and without puromycin for two weeks before proceeding with downstream analyses.

A similar procedure was used to transduce KPC cells with the lentiCRISPRv2 lentiviral vectors used for validation experiments.

### Mouse experiments

To generate pancreatic orthotopic tumors, KPC cancer cells (50’000) were orthotopically grafted into the pancreas of 8-10 weeks old syngeneic Cas9 expressing mice, or Rag2 -/- mice, obtained from the Jackson Laboratory (strains #028239 and #008449 respectively), following established protocols^51^.

For immune checkpoint blockade, ten days after injection, mice were randomized into two groups. Randomized mice were injected intraperitoneally with 200 μg per dose of InVivoMAb anti-mouse PD1 (CD279) Clone 29F.1A12 (Bioxcell #BE0273) or InVivoMAb rat IgG2a isotype control (Bioxcell #BE0089) antibodies, every third day.

For macrophage depletion experiments, mice were divided into two groups: control and macrophage depletion. Control group mice received a single dose of 1 mg IgG (InVivoMAb rat IgG1 isotype control, anti-trinitrophenol, #BE020, Bioxcell) on day 1, followed by 200 µl of control liposomes (PBS) (SKU# CLD-8914, Encapsula) on day 2. In the macrophage depletion group, mice were treated with 1 mg of InVivoMAb anti-mouse CSF1 (anti-CSF1, #BE0204, Bioxcell) on day 1 and 200 µl of clodronate liposomes (SKU# CLD-8914, Encapsula) on day 2. After one week, mice underwent orthotopic transplantation with KPC control, *Serpinb2* KO, or *Serpine1* KO cells. Next, four additional treatment cycles were administered every third day, with each cycle consisting of 0.5 mg of IgG or anti-CSF1 on the first day, followed by 200 µl of control or clodronate liposomes on the second day for the control and macrophage depletion groups, respectively. The experiment concluded after two weeks from tumor cell injection.

Animals were euthanized either upon reaching the humane endpoint for survival analyses or at the specified time points outlined in the results section and figure legends for phenotypic analyses.

### Flow cytometry

For cell sorting experiments, adherent cells were detached with 0.05% trypsin-EDTA, washed in PBS and resuspended in cell culture media. Samples were sorted on the BD FACSAria III Sorter (BD Biosciences).

For flow cytometry profiling of the TME, fresh PDAC samples were minced and enzymatically digested with the tumor dissociation kit (Miltenyi, catalog no. 130-096-730) for 40 minutes at 37°C with agitation. The cell suspension was strained through a 70 µm strainer, spun down and resuspended in flow cytometry buffer (PBS, 2% bovine serum albumin, 5 mM EDTA). Cells were centrifuged at 350 g for 5 minutes at 4°C and then pellets were resuspended with ACK lysis buffer (Life Technologies) to lyse red blood cells at room temperature for 10 min and washed with cold flow buffer. Samples were then resuspended in flow cytometry buffer and stained for 30 minutes at 4°C. All antibodies used are detailed in the **supplementary table 3**. Upon staining, cells were analyzed using a BD LSR Fortessa. Flow cytometry data were acquired using the FACS Diva software v.7 (BD) and were analyzed using FlowJo.

### CyTOF Mass cytometry

Cell suspension processing and CyTOF analysis were carried out as described previously^52^. In brief, 3×10^6^ cells were harvested, resuspended in PBS and stained for viability using Cell-ID Intercalator-103 Rh for 15 minutes at 37°C. Surface marker staining was then performed in flow buffer with an anti-mouse CD16/CD32 blocking antibody (eBioscience) on ice for 30 minutes. Cells were subsequently fixed and permeabilized using the eBioscience FOXP3/Transcription Factor Staining Buffer Set (Invitrogen) following the manufacturer’s instructions. Afterward, cells were stained with epitope-tag antibodies on ice for 1 hour and incubated with 125 nM Ir intercalator (Fluidigm) diluted in PBS with 2.4% formaldehyde at room temperature for 30 minutes. Following staining, cells were washed and stored in 10% DMSO FBS at −80°C until acquisition. Samples were acquired using either a CyTOF2 or Helios instrument (both from Fluidigm) at an event rate of <500 events per second. Antibodies were purchased in purified form and conjugated in-house using MaxPar X8 Polymer Kits (Fluidigm) according to the manufacturer’s protocol. Details of the antibodies used for mass cytometry can be found in **supplementary table 3**.

### CyTOF data analysis

CyTOF data analysis was conducted as previously detailed^52^. In summary, manual gating was performed on Cytobank to select single, live, and PC-positive (mCherry+) cells. PC-positive cells were subsequently debarcoded using the Single Cell Debarcoder tool^53^.

### Western blot

After cell culture, the medium was removed, and the dishes were washed twice with ice-cold PBS. Cells were lysed using 150 μl of RIPA buffer (Thermo Scientific, #89900) containing protease and phosphatase inhibitors. Lysis and lysate collection were carried out on ice. The lysates were incubated on ice for 5 to 10 minutes before being transferred to 1.5 ml Eppendorf tubes and centrifuged at 18,000g for 10 minutes at 4°C to collect the supernatant. Protein concentrations were quantified using the Pierce BCA Protein Assay Kit (Thermo Scientific, #23227) as per the manufacturer’s protocol. For electrophoresis, 30 µg of protein from each sample was loaded per lane along with the PageRuler™ Plus Prestained Protein Ladder (Thermo Scientific, #26619). Proteins were separated on an Invitrogen NuPAGE 10% Bis-Tris gel (Thermo Scientific, NP0315BOX) at 100 V. Subsequently, proteins were transferred onto methanol-activated PVDF membranes at 300 mA for 2 hours. The membranes were blocked in 5% milk dissolved in TBS-T, tris-buffered saline (Fisher Bioreagents, BP24711), with 0.1% Tween-20 (Fisher Bioreagents, BP337) for 1 hour at room temperature on a plate rocker. After blocking, membranes were washed three times with TBS-T (10 minutes per wash) and incubated overnight at 4°C with the indicated primary antibodies. The antibodies were diluted in the blocking buffer according to the manufacturer’s instructions. The following day, primary antibodies were removed, and membranes were washed three times with TBS-T (5 minutes per wash) on a plate rocker. Membranes were then incubated with secondary antibodies diluted 1:10,000 in TBS-T for 1 hour at room temperature with gentle rocking. Following secondary antibody incubation, membranes were washed three more times with TBS-T (10 minutes per wash) and incubated with chemiluminescence reagent (Pierce™ ECL Western Blotting Substrate, Thermo Scientific, #32209) per the manufacturer’s instructions. The following primary antibodies were used at a 1:1,000 dilution: anti-PAI1 (Invitrogen, MA1-40224), anti-PAI2 (Invitrogen, PA5-27857), and anti-Vinculin (Sigma-Aldrich, V4505). Secondary antibodies included anti-rabbit-HRP (Cell Signaling, 7074S) and anti-mouse-HRP (Cell Signaling, 7076P2).

### Multiplexed immunohistochemical consecutive staining on single slide (MICSSS)

MICSSS was carried out following a previously described protocol^21^. Briefly, 5 µm-thick FFPE tissue sections were baked at 60°C overnight, deparaffinized using xylene, and gradually rehydrated through a series of ethanol solutions (100%, 90%, 70%, and 50% in water). Antigen retrieval was achieved by incubating the slides in Antigen Retrieval Solution (pH 9, Dako) at 95°C for 30 minutes. The slides were then cooled to room temperature (RT) for 30 minutes, rinsed with TBS, and treated with 3% hydrogen peroxide at RT for 15 minutes to inhibit endogenous peroxidase activity. Following this, slides were blocked with Serum-Free Protein Block (Dako) for 30 minutes at RT and incubated with primary antibodies diluted in Antibody Diluent, Background Reducing (Dako) for 1 hour at RT. After washing with TBS containing 0.04% Tween 20, HRP-conjugated secondary antibodies were applied for 30 minutes based on the species of the primary antibody, including EnVision+ System-HRP Labelled Polymer Anti-mouse (Dako), EnVision+ System-HRP Labelled Polymer Anti-rabbit (Dako), VisUCyte HRP Polymer Goat IgG Antibody (R&D Systems), or ImmPRESS HRP Anti-Rat IgG, Mouse Absorbed (Vector Laboratories). Antigen detection was carried out using the AEC Peroxidase Substrate Kit (Vector Laboratories), and counterstaining was achieved with Harris Modified Hematoxylin Solution (Sigma-Aldrich). The slides were then mounted with Glycergel Mounting Media (Agilent) and scanned at 20x magnification using the Aperio AT2 slide scanner (Leica). For additional staining rounds, the coverslips were removed by immersing the slides in 60°C water. Residual AEC and Hematoxylin were stripped using a sequential treatment with ethanol solutions [50%, 70% (containing 1% HCl 12N), and 100%]. Slides were then processed according to the original protocol with one modification: an additional blocking step was introduced. Depending on the species of the primary antibody used for that round, the slides were incubated for 30 minutes at RT with one of the following Fab fragment reagents:

AffiniPure Fab Fragment Donkey Anti-Mouse IgG (H+L), AffiniPure Fab Fragment Donkey Anti-Rabbit IgG (H+L), AffiniPure Fab Fragment Donkey Anti-Goat IgG (H+L), or AffiniPure Fab Fragment Donkey Anti-Rat IgG (H+L) (Jackson ImmunoResearch). The antibodies utilized for MICSSS are detailed in **supplementary table 3**.

### MICSSS image processing

The MICSSS data was processed and analyzed using Fiji and QuPath, following a previously specified protocol^21^. In brief, sequentially acquired images from each staining round were aligned using the Linear Stack Alignment tool with SIFT registration in Fiji. For each image, the hematoxylin and marker staining signals were separated through deconvolution in Fiji, utilizing the default Hematoxylin/AEC color vector. Stacked layers of AEC signals for each staining round, along with one hematoxylin image, were pseudocolored and combined to generate composite images. Cell segmentation was performed in QuPath using nuclear detection on the hematoxylin stain with optimized settings. The same segmentation parameters were applied consistently across all images within the experiment.

### Pro-code debarcoding on MICSSS data

PCs were assigned to cells using a modified algorithm based on Zunder et al.^53^, as previously described in detail^12^. In summary, the mean pixel intensity for each epitope tag within the nuclear regions of segmented cells was normalized to a scale of 0 to 1 for each tissue section. Epitope tags were then ranked by their normalized intensity for each cell, and the difference between the 3rd and 4th highest intensity tags, referred to as the delta value, was calculated. Cells were assigned a PC corresponding to the three highest-intensity tags if their delta value exceeded 0.01. These PC assignments were cross-referenced with the vector library design to identify the associated sgRNA target genes. PC assignments were further validated by applying marker-specific intensity thresholds, which were predefined for each epitope based on optimization across tissue sections. Cells that did not meet the intensity thresholds for all three markers within a PC code combination were excluded from the final assignment. All images were then combined in a large single cell object and further analyzed using Squidpy^54^ and visualized in R (v4.2.2).

### Enrichment and depletion analysis

To assess enrichment or depletion relative to the internal control of the library, we first calculated the number of PC+ cells for each mouse. The PC+ cell counts were normalized against the total number of PC+ cells within each mouse to account for variations in overall cell numbers. Individual animals were treated as biological replicates to ensure statistically robust analysis. We applied multiple Mann-Whitney tests to compare each condition against the control, using the null hypothesis that no significant differences existed.

### Tumor clonality assessment on MICSSS data

To assess tumor clonality on MICSSS data, we analyzed spatially resolved cell coordinates of PC+ cells. We constructed k-nearest neighbor (kNN) graphs to quantify clonal purity by calculating the fraction of neighboring cells with mismatched barcodes. Dense clonal regions, or focal areas, were identified using mean shift clustering with a 250 μm bandwidth, retaining clusters with at least 50 cells. Clonal diversity was quantified within these focal points using the Shannon Diversity Index (SDI) and the evenness index, capturing both heterogeneity and uniformity. Temporal and spatial variations in clonality were analyzed by calculating median SDI and evenness across conditions, highlighting dynamic changes over time. Spatial clonality patterns were visualized with spatial plots, overlaid with focal point polygons, while stacked bar plots illustrated barcode frequencies within clusters. All analyses were performed using R (v4.2.2).

### Neighborhood enrichment analysis

To investigate spatial relationships between cell clusters across tumor tissues, we performed neighborhood enrichment analysis using Squidpy (v1.3.0). The neighborhood enrichment scores were computed using the squidpy.gr.nhood_enrichment function, which employs a permutation-based test. Clusters that frequently co-occur as neighbors across the tissue exhibit higher scores, indicating spatial enrichment. Conversely, clusters with low scores are spatially segregated, suggesting depletion. We used 10,000 permutations to ensure robust statistical assessment, unless otherwise specified. The enrichment scores (Z-scores), for pro-code analysis and Perturb-map at days 7, 14 and 21, were extracted and saved for further interpretation and visualization.

### MICSSS spatial distances

Spatial distance analysis between specific phenotypes was performed using the squidpy.tl.var_by_distance function in Squidpy (v1.3.0). For each cell of a particular phenotype within the defined ROIs, the shortest distance to a designated anchor point was calculated at the single-cell level. The resulting distances were visualized as distribution plots in the corresponding figure panels. Additionally, median distances were computed for each ROI and each tumor/mouse and represented as violin plots.

### Masson’s trichrome staining and analysis

Masson’s trichrome staining was carried out using Trichrome Stain Kit (Connective Tissue Stain, Abcam # ab150686) on 5 µm-thick FFPE tissue sections following the manufacturer’s instructions.

For data analysis, tumor images were processed in Fiji. The Colour Deconvolution2 plugin with the “Masson Trichrome” vector was used to isolate collagen-specific staining. Area measurements were performed to quantify collagen content across all samples in a batch-wise manner.

### DepMap analyses

For the visualization of DepMap data, we utilized the CRISPR (DepMap Public 22Q4+Score, Chronos) dataset, which was downloaded from the DepMap portal (https://depmap.org/portal/)^55^. This dataset included 45 human PDAC cell lines. In this dataset, a dependency score of 0 indicates that the gene is not essential for cell viability, while a score of -1 represents the median dependency score of universally essential genes, providing a benchmark for assessing gene essentiality.

### Pan Cancer bulk RNA-seq analysis

Data used for the Pan Cancer bulk RNA-seq analysis was generated by the TCGA (https://www.cancer.gov/tcga) and GTEx (https://www.gtexportal.org) projects. The raw data from all three projects was re-processed by Vivian et al., in an effort to reduced batch-effects between the datasets^56^. In the same processing the data was log2-normalized. This re-processed data was accessed from the UCSC Xena Data Hubs via the UCSCXenaTools R package^57^. Specifically, the following datasets were used for the study: TcgaTargetGtex_RSEM_Hugo_norm_count, TcgaTargetGTEX_phenotype, TCGA_survival_data. Differences in normalized expression levels of SERPINE1 and SERPINB2 between cancer subtypes and normal tissue were tested using a two-sided Wilcoxon rank sum test. Subsequently, p-values were adjusted for multiple testing by the Benjamini-Hochberg correction. Patients in each cancer type were stratified into high- and low-expression groups for both genes using median gene expression as the threshold. To assess the survival advantage in the low-expression groups, the *coxph* function of the survival package (v3.7.0) was employed to calculate hazard ratios and p-values. To analyze survival specifically in PDAC within the TCGA dataset, the data was subset, and Kaplan-Meier survival curves were generated using the *survfit* function of the survival package (v3.7.0), with p-values calculated via log-rank tests.

### Survival Analysis in Patients Treated with Immunotherapy

Bulk RNA-seq data and clinical data from Motzer et al. were analyzed to assess the impact of SERPINE1 and SERPINB2 expression on progression-free survival (PFS) in patients treated with Avelumab plus Axitinib^38^. Patients were stratified into high- and low-expression groups for both genes using median gene expression as the threshold. Kaplan-Meier survival curves were fitted using the *survfit* function of the survival package (v3.7.0) and p-values were computed with a log-rank test.

### Bulk RNA-seq

Control, *Serpinb2* and *Serpine1* KO cells, were seeded in 10 cm dishes. The following day, they were detached and resuspended in 1X PBS. The PBS was removed, and cell pellets were stored at -80°C until further use. RNA isolation, sample QC, library preparation with poly(A) selection and 150-bp paired-end sequencing (30 million reads/sample) were performed by GENEWIZ (South Plainfield, NJ).

### Bulk RNA-seq analyses

Paired-end FASTQ files were evaluated for quality using FastQC (v0.12.1), and trimmed with cutadapt (v4.9). Trimmed reads were then aligned to the GRCm39 mouse assembly with the corresponding GENCODE Release 36 gene annotation using STAR (v2.7.11b). Normalized counts were calculated with the estimated size factors within DESeq2 (v1.44.0), and transformed with variance-stabilizing transformation. The Principal Component Analysis (PCA) was carried out on the transformed counts with PCAtools (v2.16.0), after removing the lowest 10% of variance. Differential expression testing was performed between the knockout and control samples with the design corresponding to the KOs (‘∼KÒ), with results extracted individually for each KO (*Serpine1* KO and *Serpinb2* KO) against the control. Subsequent enrichment analyses were performed using the Enrichr database^58^, with cutoffs used for the specific analyses being specified in the text and corresponding figure legends.

### Mouse single cell RNA sequencing (scRNA-seq)

Fresh tumor samples were processed into single-cell suspensions following the “Flow Cytometry” tumor processing protocol described earlier. Cell viability was assessed using the Acridine Orange/Propidium Iodide viability staining reagent (Nexcelom). Suspensions with over 80% viability and minimal debris were deemed suitable for downstream experiments. scRNA-Seq was conducted using the Chromium platform (10x Genomics) with the 5’ Gene Expression (5’ GEX) V2 kit, with a targeted recovery of 8,000 cells. Gel-Bead in Emulsions (GEMs) were generated on the sample chip K using the Chromium X (10x Genomics). Barcoded cDNA was extracted from GEMs through Post-GEM RT cleanup and amplified via 13 PCR cycles. Amplified cDNA was fragmented, end-repaired, poly A-tailed, adapter-ligated, and sample-indexed according to the manufacturer’s instructions. Libraries were quantified using TapeStation (Agilent) and QuBit (ThermoFisher) and sequenced in paired-end mode on an Illumina NovaSeq 6000 instrument, with a target depth of 25,000 reads per cell. Raw FASTQ files were aligned to the reference genome Gex-mm10-2020-A using CellRanger v5.0.1 (10x Genomics). Feature barcode matrix files generated by CellRanger were utilized for downstream analyses.

### scRNA-seq analyses

ScRNA-seq data were processed using the Scanpy library (v1.9.4). Data from the Control, *Serpinb2* KO, and *Serpine1* KO conditions were concatenated into a single AnnData object for joint analysis. Quality control included filtering cells with fewer than 800 or more than 40,000 total counts. Cells with mitochondrial gene content exceeding 20% of total counts were excluded to account for potential stress or degradation. Additionally, cells expressing fewer than 300 genes were removed. Doublets were identified using Scrublet (v0.2.3) with default parameters and filtered based on a threshold doublet score of 0.04. After filtering, 15,815 cells were retained for downstream analysis. Genes with fewer than 20 total counts across all cells were removed, leaving 16,380 genes for analysis. Raw counts were normalized using a log-transformation with a pseudocount of 1 to stabilize variance across cells. Highly variable genes were identified using Scanpy’s highly_variable_genes function, focusing on the top 4,000 genes based on dispersion and mean expression across cells. The processed dataset was used for subsequent normalization, dimensionality reduction, clustering, and annotation.

Differential gene expression analysis was performed with the tool rank_genes_groups, which is part of the Scanpy package. The Benjamini–Hochberg method was used to correct for multiple testing. Subsequent enrichment analyses were performed using the Enrichr database^58^, with cutoffs used for the specific analyses being specified in the text and corresponding figure legends.

The fibrinogen binding score was calculated to evaluate the enrichment of fibrinogen-associated gene signatures in macrophage subpopulations. A curated list of fibrinogen-binding genes, derived from the “fibrinogen binding” molecular function Gene Ontology (GO) term, was used as the basis for this analysis. The score_genes function in Scanpy was employed to calculate the scores, reflecting the expression levels of this gene set across the macrophage cluster.

The additional dataset analyzed can be found at GSE207938.

### Human scRNA-seq

The two analyzed dataset can be found under the GEO accession number GSE155698 and via the Human Tumor Atlas Network (HTAN) dbGaP Study Accession phs002371.v1.p1. First, barcodes from the processed Zhou et al. scRNA-seq dataset were filtered after calculating quality control variates across all samples. Potential empty droplets were removed if the total transcript counts were too few (<100), if too few genes were expressed (<150), or if the total transcript counts were too high (>40,000). Barcodes with a mitochondrial fraction of over 20% were removed. Second, Scrublet was used to determine a doublet filter of 0.14, with barcodes removed if they fell above this threshold. Finally, batch correction across samples was conducted with Harmony’s integrate function, where the samples converged after 3 iterations. Cells in the final AnnData (v0.10.9) object were clustered using leiden clustering with Scanpy’s (v1.10.0) ‘sc.tl.leiden’ methodology, using ‘scanpy.pp.neighbors’ with 7 PCA dimensions and ‘scanpy.t.umap’.

### Human scRNA-seq analysis

Differential expression analysis was carried out using scanpy.tl.rank_gene_groups. SERPINE1 and SERPINB2 cancer cells were compared to all other cancer cells with the Wilcoxon method. Significantly differentially expressed genes (p < 0.05) that were upregulated and downregulated were separately used for the enrichment analysis. Enrichment analyses were performed using the Enrichr database^58^, with cutoffs used for the specific analyses being specified in the text and corresponding figure legends.

Cell type annotation was conducted through manual evaluation of defined key marker genes in each cluster, using a mixture of known markers and original markers used in the Zhou et al. scRNA-seq dataset. Cancer cells were further classified using a “Collison Score” based on predefined marker genes for Classical, Basal, and Exocrine-Like states. The mean expression of these markers was calculated for each cluster and state, followed by the median variance across clusters. A cluster was assigned to a specific Collison state if its marker gene variance was higher than that of other clusters. The following genes were used: Classical (TMEM45B, SDR16C5, GPRC5A, AGR2, S100P, FXYD3, ST6GALNAC1, CEACAM5, CEACAM6, TFF1, TFF3, CAPN8, FOXQ1, ELF3, ERBB3, TSPAN8, TOX3, LGALS4, PLS1, GPX2, ATP10B, MUC13), Basal (AIM2, GPM6B, S100A2, KRT14, CAV1, LOX, SLC2A3, TWIST1, PAPPA, NT5E, CKS2, HMMR, SLC5A3, PMAIP1, PHLDA1, SLC16A1, FERMT1, HK2, AHNAK2), and Exocrine-Like (REG1B, REG3A, REG1A, CEL, PNLIP, PLA2G1B, CELA3A, CPB1, CELA3B, CTRB2, CLPS, CELA2B, PRSS2, PRSS1, GP2, SLC3A1, CFTR, SLC4A4, SPINK1).

Cancer cells were additionally stratified by SERPINE1/B2 status. Barcodes that contained transcripts for SERPINE1, but not SERPINB2, were annotated as ‘SERPINE1’. Barcodes that only contained transcripts for SERPINB2, and not SERPINE1, were annotated as ‘SERPINB2’. Barcodes that expressed both were annotated ‘SERPINE1/B2’, and barcodes that were negative for both were annotated ‘Other’.

### Human spatial transcriptomics

Individual raw AnnData files for each 10X Visium Spatial Transcriptomic sample were downloaded from the Human Tumor Atlas Network (HTAN). Scanpy (v1.10.1) was used to calculate quality metrics. Data were integrated and normalized using the standard Scanorama pipeline. Briefly, samples were individually normalized, and log-transformed, and highly variable genes (n=2000) were calculated individually using the Seurat method. Data was corrected using scanorama_correct, and subsequently concatenated.

### Spatial transcriptomic deconvolution and cell-type specific gene expression

Spatial transcriptomic data was deconvoluted using cell2location where single-cell reference data from the same study was used to estimate reference cell type signatures. Genes were chosen from the single-cell data using cell_count_cutoff = 10, cell_percentage_cutoff2 = 0.03, and nonz_mean_cutoff = 1.12. The regression model was trained using sample ID as a batch key, treatment as a categorical covariate, and annotated cell types. The model was trained with maximum epochs being 250, and a batch size of 2500. The estimated gene expression of every cell type was exported, and subsequently used to map cell types in each spatial spot in the spatial transcriptomic data. The expected expression of each gene in each cell-type was then calculated using the posterior distribution as per the cell2location’s standard vignette. Gene expression of individual genes was plotted in spatial coordinates using cell2location’s custom plot_genes_per_cell_type function, obtained from GitHub.

### Spatial tumor microenvironment analysis of SERPINE1+ and SERPINB2+ spots

Visium spots positive for tumor cells were defined as spots containing more than 4 predicted tumor cells, as per cell2location’s posterior distribution. These cells were further stratified by SERPINE1 and SERPINB2 expression. A fixed radius of 10 around each SERPINE1+ or SERPINB2+ tumor spot was defined using the cdist function from scipy.spatial.distance. Cells within this radius were aggregated to compute cell-type abundances, and distances between cells and tumor spots were calculated using spatial coordinates from the dataset. Cell-type counts were normalized to the total number of cells within the radius, resulting in proportional values that represent the relative abundance of each cell type.

## Statistical analysis

Statistical analysis was conducted using GraphPad Prism 10 or using the software indicated above. Details of statistical analysis are described in the figure legends.

## Materials availability

The nuclear Pro-Code library is available from Addgene (Plasmid Kit #1000000197). All other vector constructs used in the manuscript are available by request to the lead contact.

## Data and code availability

The original code present in this manuscript can be found at GitHub (https://github.com/BDBrownLab). The bulk and single-cell transcriptomics data are available at the Gene Expression Omnibus (GEO) webserver under accession ID GSE289138.

## Supplementary figures

**Figure S1.**
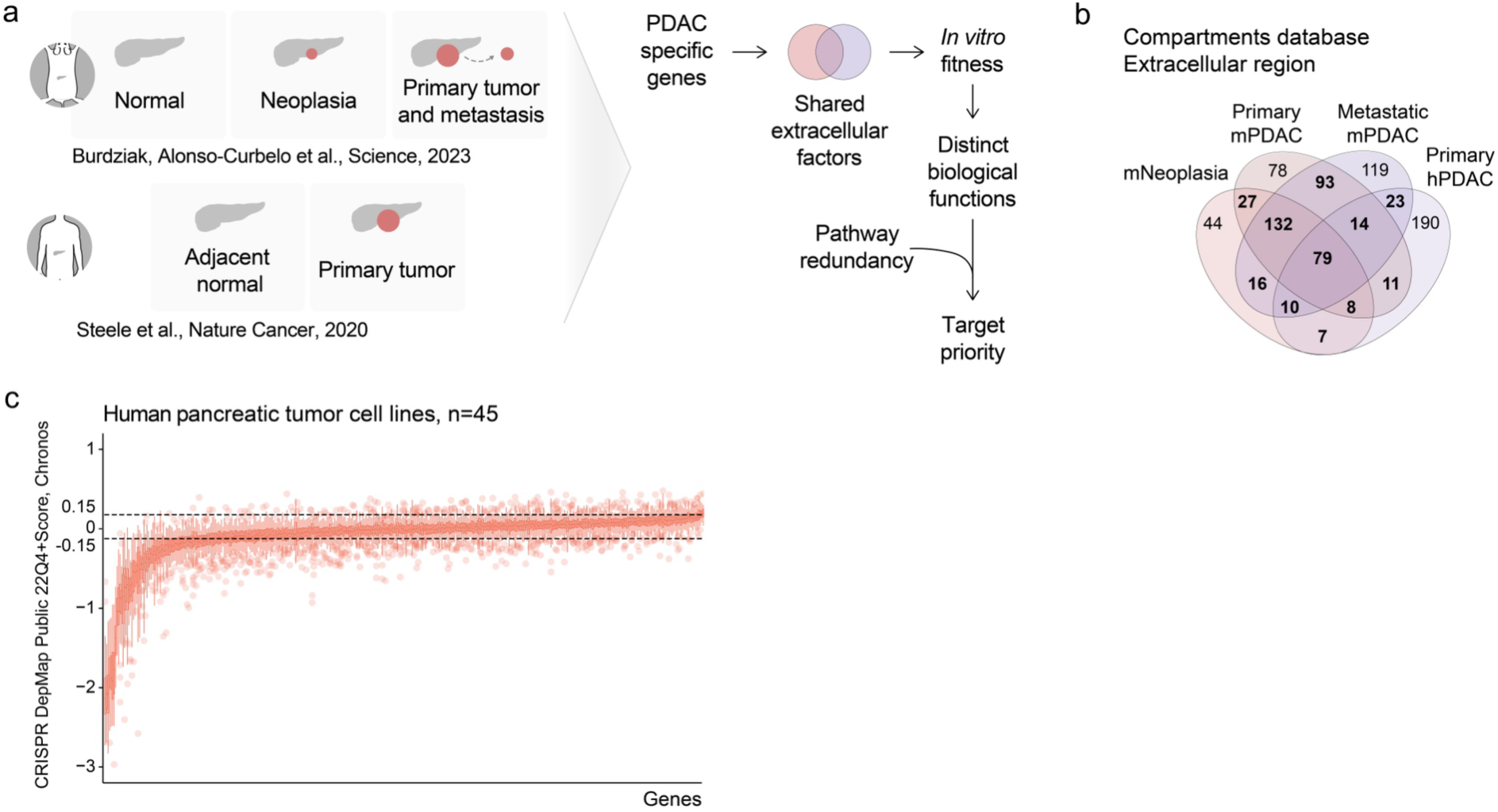
Systematic identification and prioritization of genes with extracellular functions to study tumor host interactions that drive PDAC progression. **a**, Schematic workflow illustrating the approach to identify and prioritize extracellular factors involved in shaping the PDAC TME. **b**, Differential gene expression analysis was performed to compare transformed or tumor cells with normal or adjacent normal tissues. Two scRNA-seq datasets derived from KPC mouse models and human samples^14,15^ were used. The analysis prioritized PDAC-specific genes with an adjusted p value ≤ 0.05 and a log2 fold change > 4 vs normal controls. Subsequently, an enrichment analysis utilizing the Compartments database of protein subcellular localization was performed^16^. The overlap of “extracellular region part”-associated genes across datasets is shown. **c**, CRISPR/Cas9 dependency analysis was performed on genes showing overlap from panel (**b**), using data from 45 human PDAC cell lines subjected to *in vitro* CRISPR screens. Genes showing a median viability between -0.15 and 0.15 were used for downstream analysis. Data was obtained from the DepMap 22Q4 Public release (https://depmap.org/portal/download/). The line in the box plots shows the median value. PDAC: pancreatic ductal adenocarcinoma, TME: tumor microenvironment, scRNA-seq: single cell RNA sequencing.

**Figure S2.**
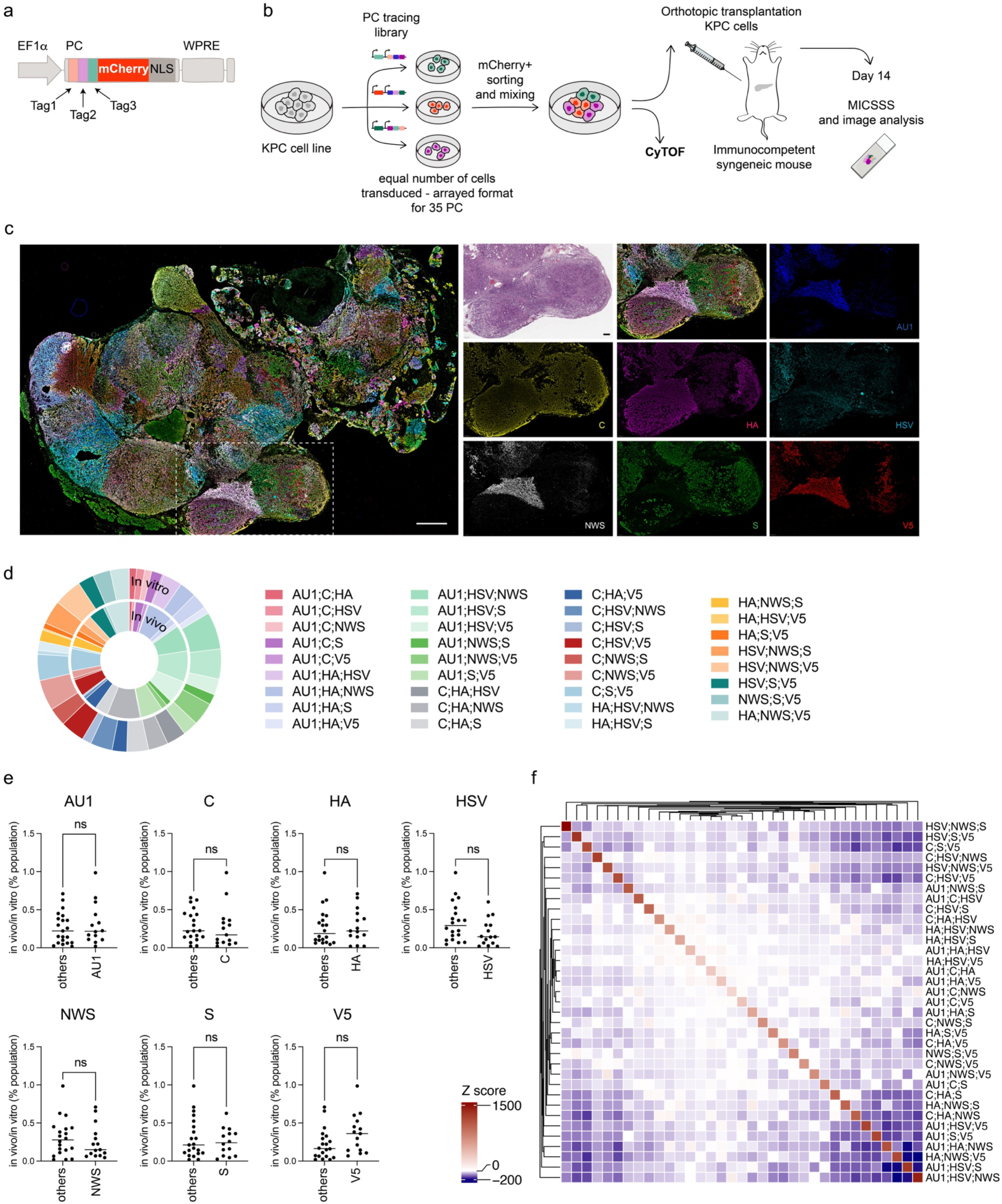
A pro-code library to investigate PDAC tumor growth. **a**, Schematic representation of the PC vector. **b**, Experimental set-up used to validate the system in PDAC. An equal number of KPC cells was seeded and transduced with the PC tracing library, composed of 35 PCs, 7 tags, in an arrayed format. Cells were sorted based on mCherry positivity and mixed in equal amounts. CyTOF was performed on the *in vitro* pool of cells prior to injection. The same cells were used for orthotopic transplantation in the pancreas of immunocompetent syngeneic mice. After 2 weeks the tumors were harvested and subjected to MICSSS analysis. One tumor section from 13 individual mice was used for analysis. **c**, Representative image of one tumor resulting from the injection of KPC cells transduced with the PC tracing library. H&E and single stains are shown on the right side. Scale bars: 1 mm for the main image and 200 μm for the insets. **d**, Pie charts showing the relative frequencies of each PC *in vitro* (outer ring) and *in vivo* (inner ring). Each PC is labeled with a distinct color. **e**, Comparisons of the relative frequencies (*in vivo*/*in vitro*) for the single epitope tags against every other tag composing the PCs. A two-tailed, unpaired Student’s t-test was performed to calculate the p values. **f**, Neighborhood enrichment analysis quantifying the interaction between single PC+ populations in PDAC tumor lesions. Each square represents an interaction between two PC populations and is color-coded based on the significance of the interaction relative to a permuted null distribution generated by swapping pro-code labels (1000 permutations). PDAC: pancreatic ductal adenocarcinoma, PC: pro-code, MICSSS: Multiplexed Immunohistochemical Consecutive Staining on Single Slide, ns: non-significant.

**Figure S3.**
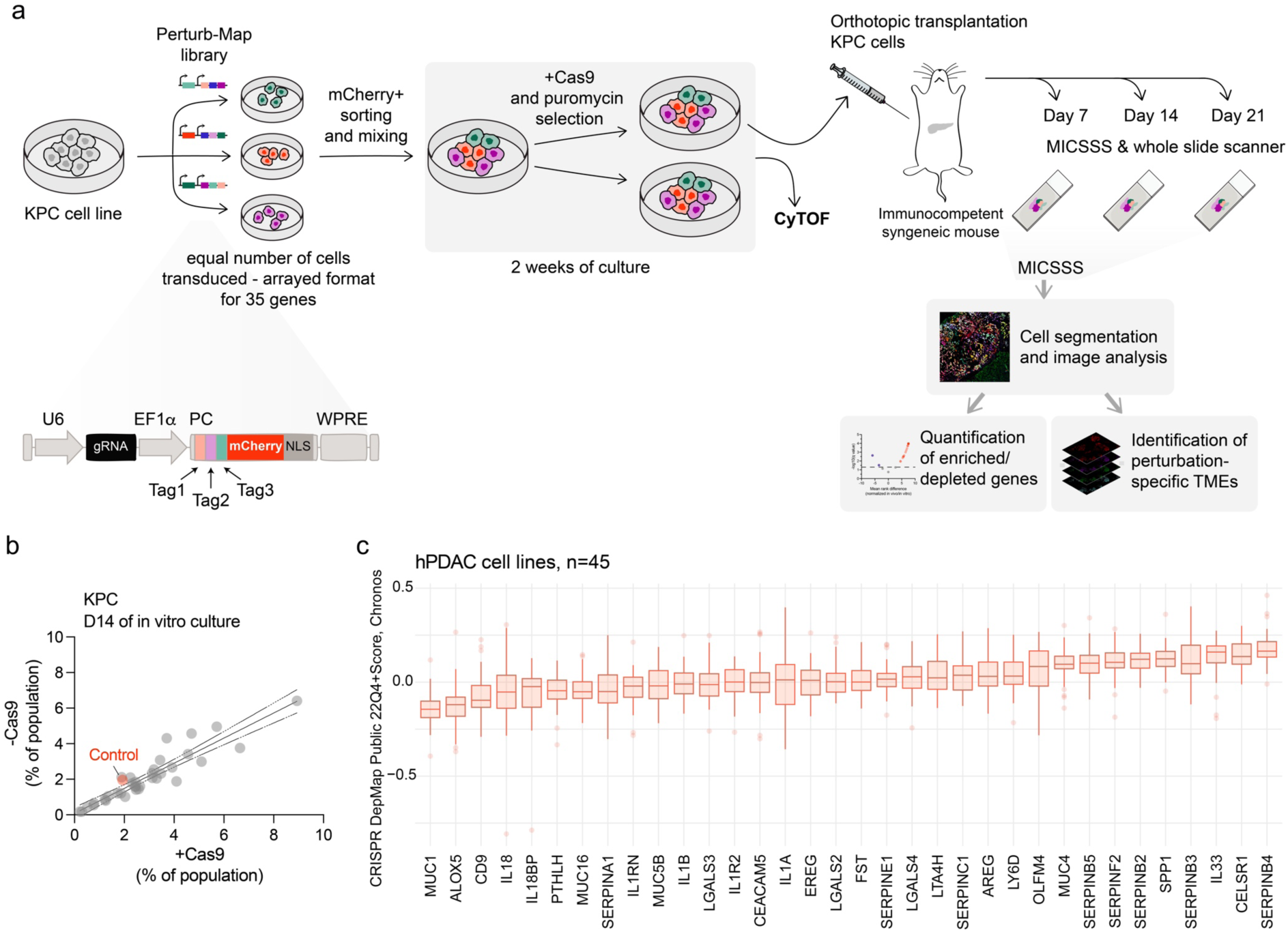
The perturbed genes do not alter PDAC growth *in vitro*. **a,** Detailed scheme of the experimental set-up to study PDAC extracellular factors in immunocompetent mice across time points and schematic representation of the Perturb-map vector used in the study. **b,** KPC cells were transduced with the Perturb-map library and divided into two arms. One arm was further transduced with Cas9 and selected with puromycin for 7 days. Both arms were then cultured for an additional 2-week period. The relative frequency of each PC in the Perturb-map library is shown, comparing conditions with (x-axis) and without (y-axis) Cas9. **c,** DepMap viability scores for the human orthologues of the selected genes across 45 human pancreatic cancer cell lines. Data was obtained from the DepMap 22Q4 Public release (https://depmap.org/portal/download/). MUC5AC was not included in the graph because screened only in two cell lines. The line in the box plots shows the median value. PDAC: pancreatic ductal adenocarcinoma, hPDAC: human PDAC, PC: pro-code, MICSSS: Multiplexed Immunohistochemical Consecutive Staining on Single Slide.

**Figure S4.**
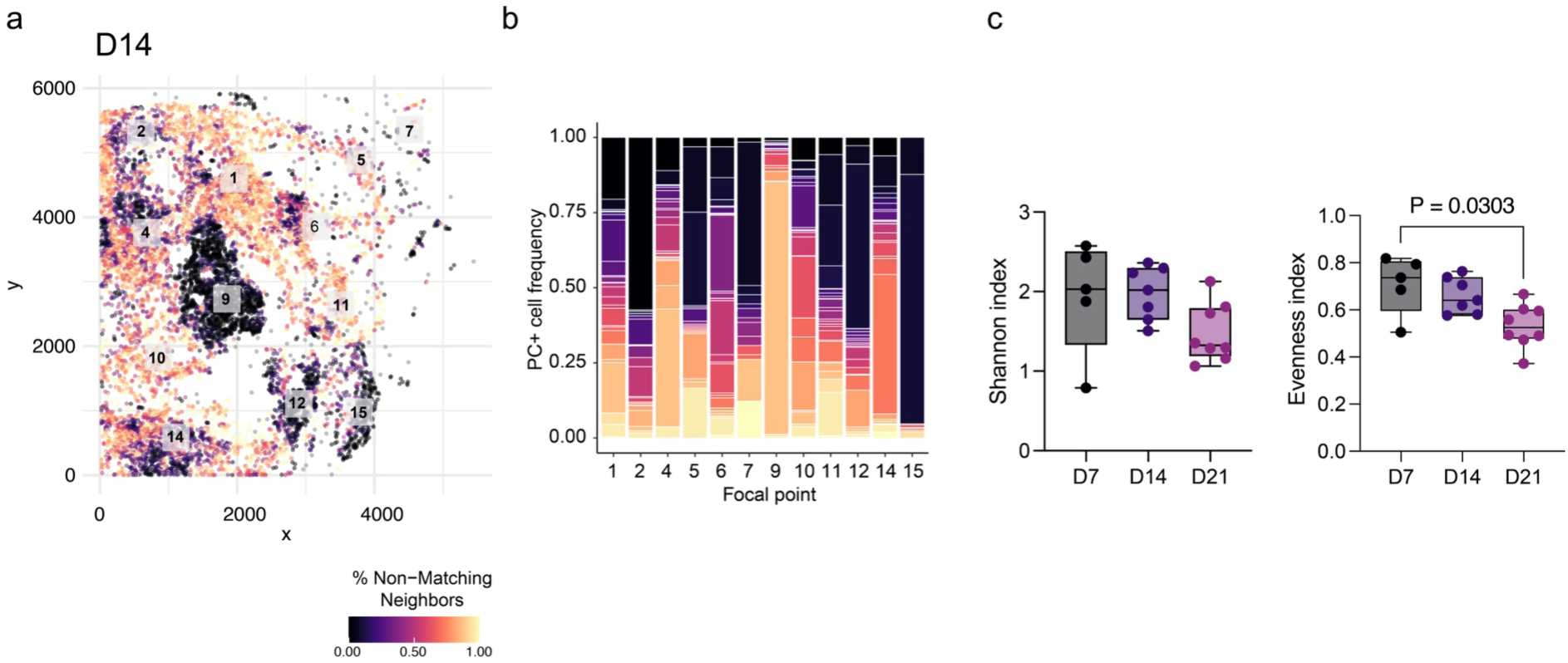
Spatial and clonal evolution of PC+ pancreatic tumor cells reveals increasing homogeneity and focal consolidation over time. **a,** Representative digital reconstruction of a tumor section isolated 14 days post-orthotopic transplantation. The color scale indicates the percentage of non-matching neighbors, where yellow represents areas with higher heterogeneity, and black represents more homogeneous regions. White squares mark focal points, identified using mean-shift clustering to detect regions of high local cell density based on a defined bandwidth. **b,** Frequency of PC+ cells within the focal areas identified in panel (**a**), highlighting the distribution of specific cell populations. PC: pro-code.

**Figure S5.**
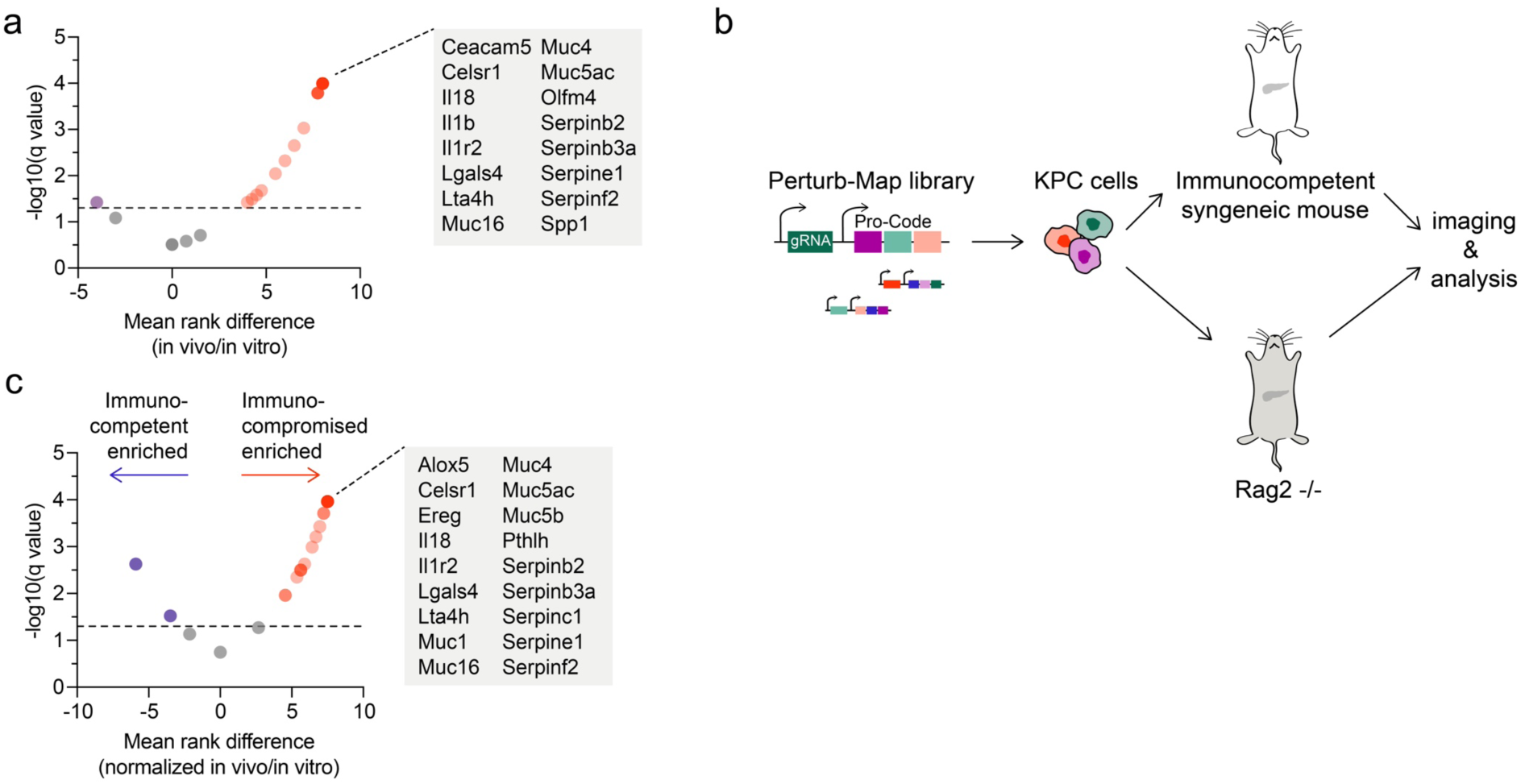
Perturb-map for identifying PDAC extracellular factors in immunocompetent and immunocompromised mice. **a**, Volcano plots comparing each gene in the library to the internal control of the screen conducted in immunodeficient Rag2-/- mice. The analysis uses the ratio of *in vivo* to *in vitro* PC counts (*i.e.*, gene KO), with significance assessed using multiple Mann-Whitney tests. Genes with an adjusted p value < 0.05 are highlighted in red (enriched) or in blue (depleted). **b**, Schematic representation of the experimental approach comparing the role of extracellular factors in immunocompetent and immunocompromised mice. **c**, Volcano plots comparing gene expression between immunocompetent and immunocompromised (Rag2-/-) mice. The immunocompetent cohort (day 14 time point) already introduced in Figure 1 was used for comparative analyses. The analysis employed the ratio of *in vivo* to *in vitro* PC (*i.e.*, gene KO) counts normalized to the internal control, with significance determined by multiple Mann-Whitney tests. Genes with an adjusted p value < 0.05 are highlighted in red (enriched in immunocompromised) or in blue (depleted in immunocompromised). PDAC: pancreatic ductal adenocarcinoma, PC: pro-code.

**Figure S6.**
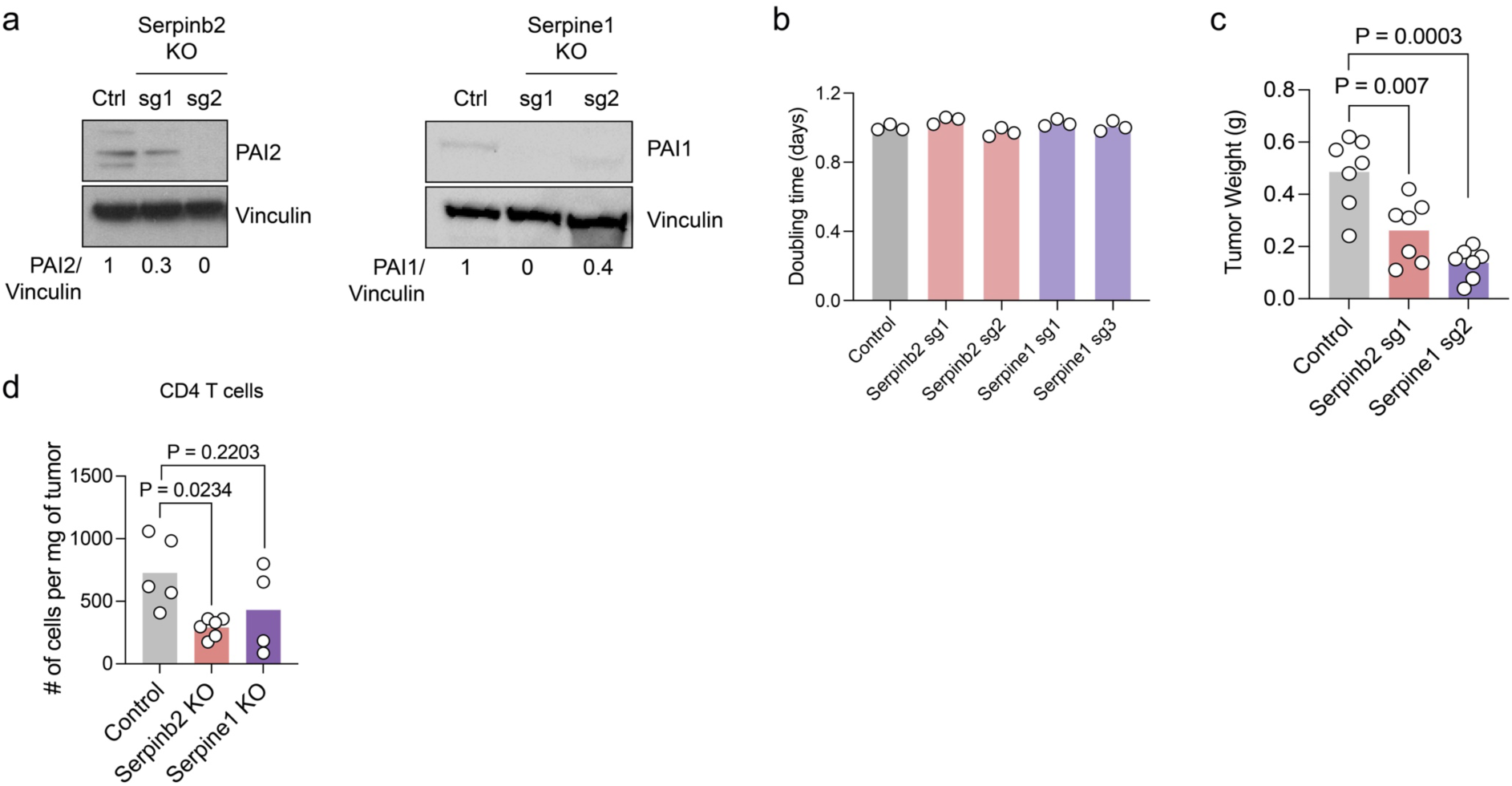
*Serpinb2* and *Serpine1* KO do not alter PDAC cell growth *in vitro*. **a**, Western blot analysis showing PAI1 (Serpine1) and PAI2 (Serpinb2) expression in KPC cell cultures with the indicated genetic perturbations using two different single guide RNAs per gene. Vinculin was used as a loading control. Protein quantification relative to the loading control is provided below each blot. **b**, Doubling time (in days) of cell lines with the indicated genetic perturbations relative to control (F8 KO) (n = 3 independent experiments). **c**, Tumor weight comparison at day 14 between control, *Serpinb2* KO (sg1), and *Serpine1* (sg2) KO tumors (control n=7, *Serpinb2* KO n=7, *Serpine1* KO n=7). The control cohort is the same shown in Figure 3c. *Serpinb2* sg2 and *Serpine1* sg1 were used to perform all additional experiments. **d**, Flow cytometry analysis of tumors isolated from control, *Serpinb2*, and *Serpine1* KO tumor bearing mice two weeks after orthotopic transplantation, showing the absolute number of CD4+ T cells (control n=5, *Serpinb2* KO n=6, *Serpine1* KO n=4). Statistical significance in **b**, **c** and **d** was determined using a two-tailed, unpaired Student’s t-test. PDAC: pancreatic ductal adenocarcinoma, KO: knock-out.

**Figure S7.**
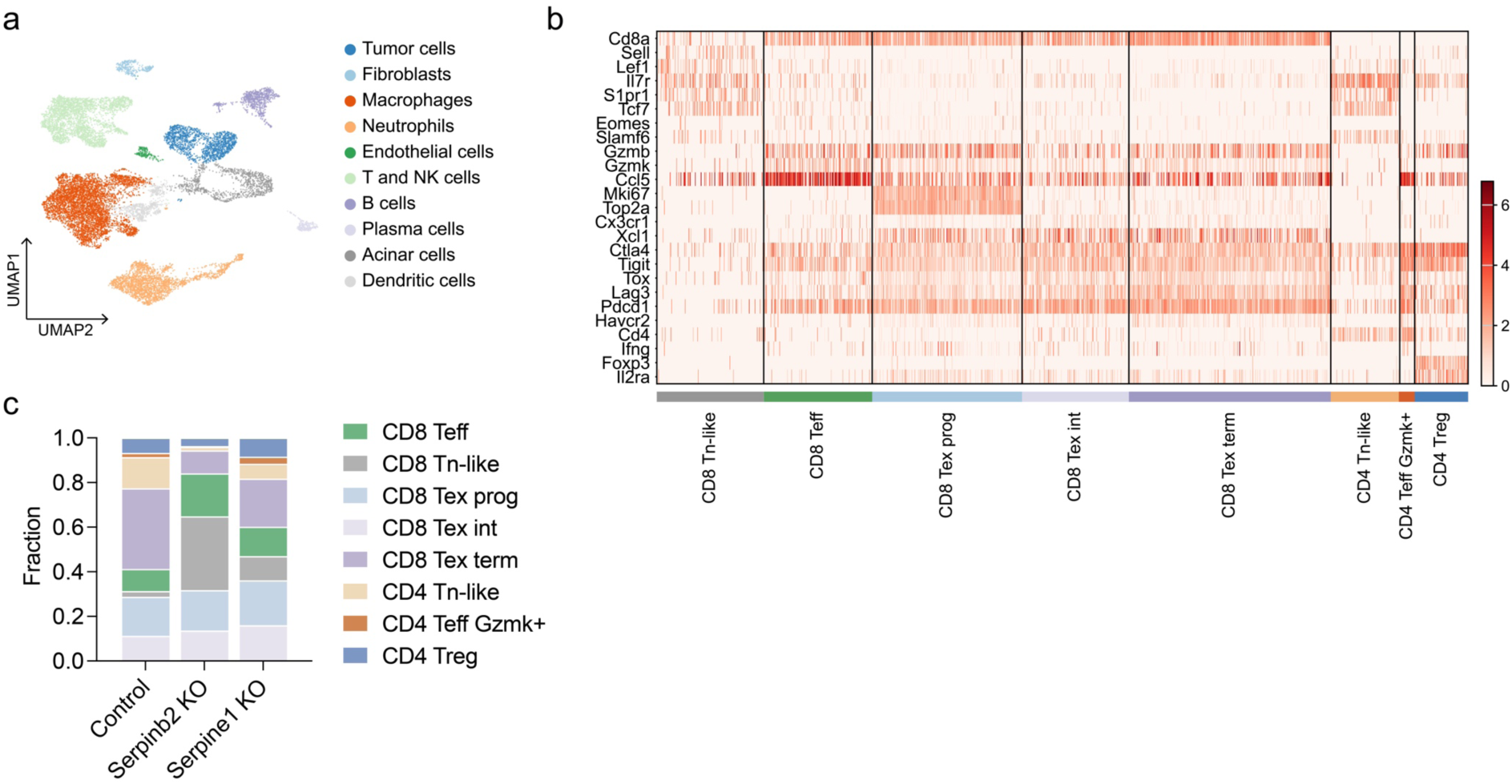
ScRNA-seq of *Serpinb2* and *Serpine1* KO tumors reveals shifts in T cell subsets that are linked to anti-tumor immunity. **a**, UMAP projection showing cell populations identified in the scRNA-seq experiment across control, *Serpinb2* KO, and *Serpine1* KO tumors. **b,** Heatmap of hallmark genes across the identified T cell subsets. Normalized gene expression is shown. **c,** Relative frequency of all identified T cell subsets across the investigated conditions. KO: knock-out, scRNA-seq: single cell RNA sequencing.

**Figure S8.**
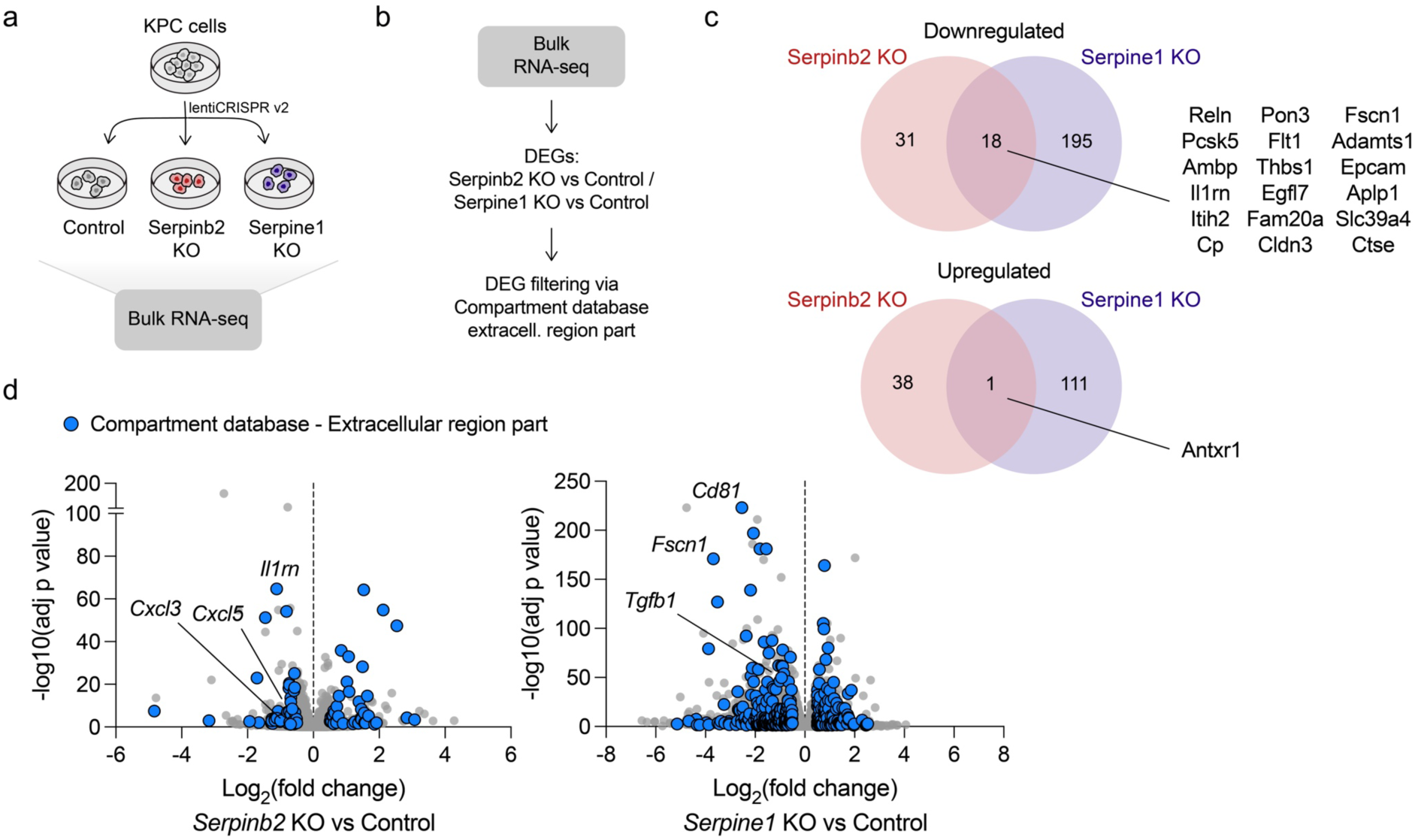
Transcriptome changes upon *Serpinb2* and *Serpine1* KO. **a**, Schematic overview of the bulk RNA-seq experiment. **b**, Workflow for identifying differentially expressed extracellular factors following *Serpinb2* and *Serpine1* KO. DEGs were identified by comparing *Serpinb2* KO and *Serpine1* KO cells to controls, with filtering criteria of adjusted p value ≤ 0.05 and absolute log2 fold change ≥ 0.5. Filtered DEGs were cross-referenced with the “extracellular region part” category in the protein subcellular localization compartments database. **c**, Venn diagram illustrating overlap between upregulated and downregulated DEGs with extracellular roles in *Serpinb2* and *Serpine1* KO cells. Shared genes across both conditions are highlighted. **d**, Volcano plots showing gene changes in *Serpinb2* KO (left) and *Serpine1* KO (right) cells compared to controls. Upregulated and downregulated genes annotated with extracellular region part for both KOs are highlighted in blue. Selected genes are labeled. KO: knock-out, RNA-seq: RNA sequencing, DEGs: differentially expressed genes.

**Figure S9.**
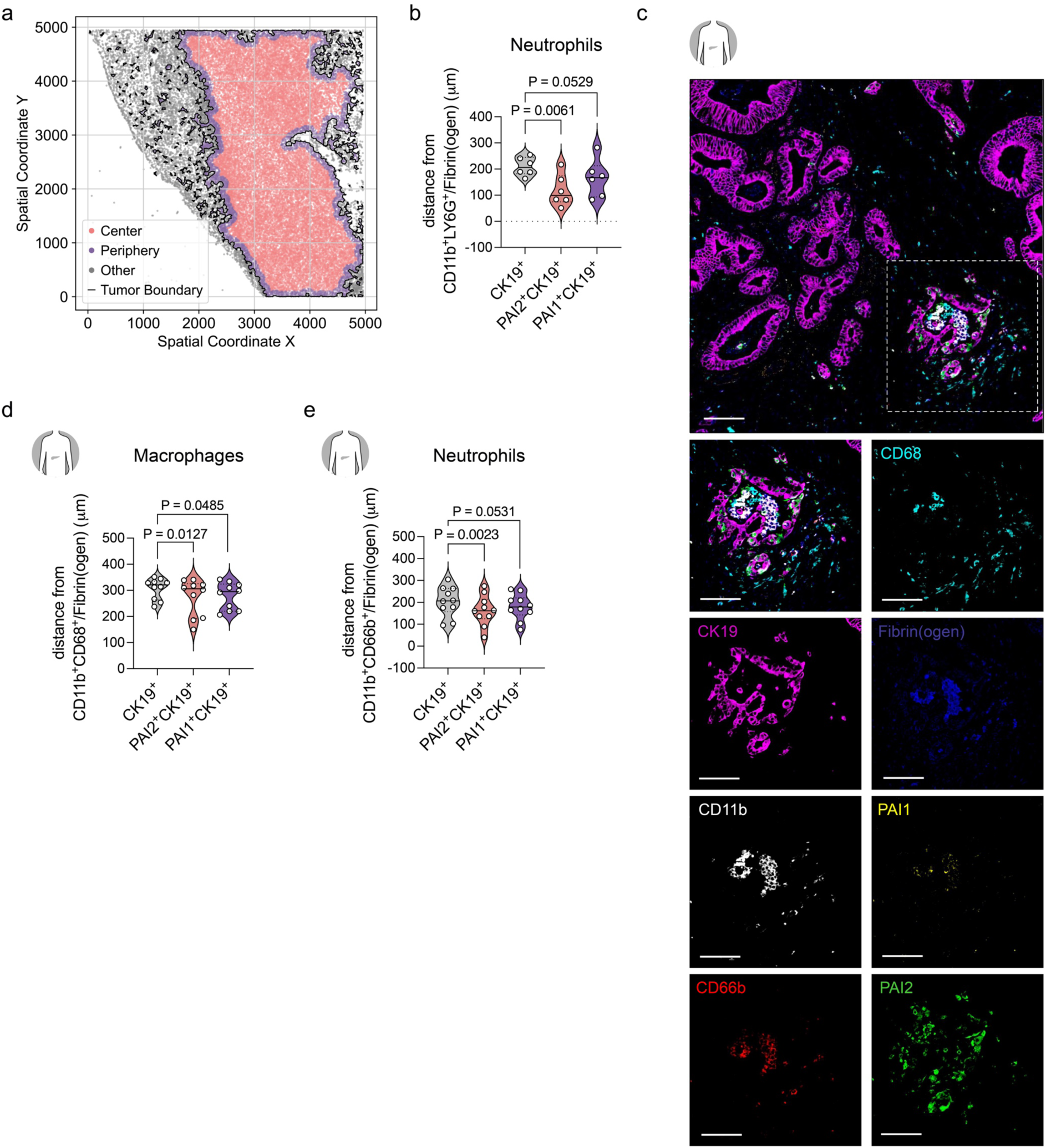
PAI1 and PAI2 expression in CK19+ tumor cells and spatial relationship with immune components. **a**, Representative digital reconstruction of tumor regions based on spatial distribution of CK19+ cells. The tumor boundary was defined by the presence of CK19+ cells. Regions within a 50 μm distance from the tumor boundary were designated as the “Periphery” (outlined in blue), while the interior region beyond this boundary was labeled as the “Center” (red). Areas outside the periphery were classified as “Other” (grey). The plot shows spatial coordinates (X and Y) across the tumor section, highlighting distinct spatial compartments used to perform the subsequent analyses on mouse tumors. **b**, Comparison of distances between CD11b+LY6G+ cells colocalizing with Fibrin(ogen) (CD11b+LY6G+/Fibrin(ogen)) cells, and CK19+, PAI1+CK19+, and PAI2+CK19+ cells in mouse control tumors (n=6). Only the tumor center area, as visualized in panel (**a**), was used for the analysis. **c**, Pseudocolored immunohistochemistry of hPDAC tumor sections stained for CK19, CD11b, CD66b, CD68, Fibrin(ogen), PAI1, and PAI2. Representative images are shown. Scale bar: 100 µm for the main image and 50 µm for insets. **d**, **e,** Comparison of distances between CD11b+CD66b+/Fibrin(ogen) cells (**d**), CD11b+CD68+/Fibrin(ogen) cells (**e**), and CK19+, PAI1+CK19+, and PAI2+CK19+ cells in hPDAC (n=10). P values in **b**, **d**, and **e** were calculated with a two-tailed, paired t-test. PDAC: pancreatic ductal adenocarcinoma, hPDAC: human PDAC.

**Figure S10.**
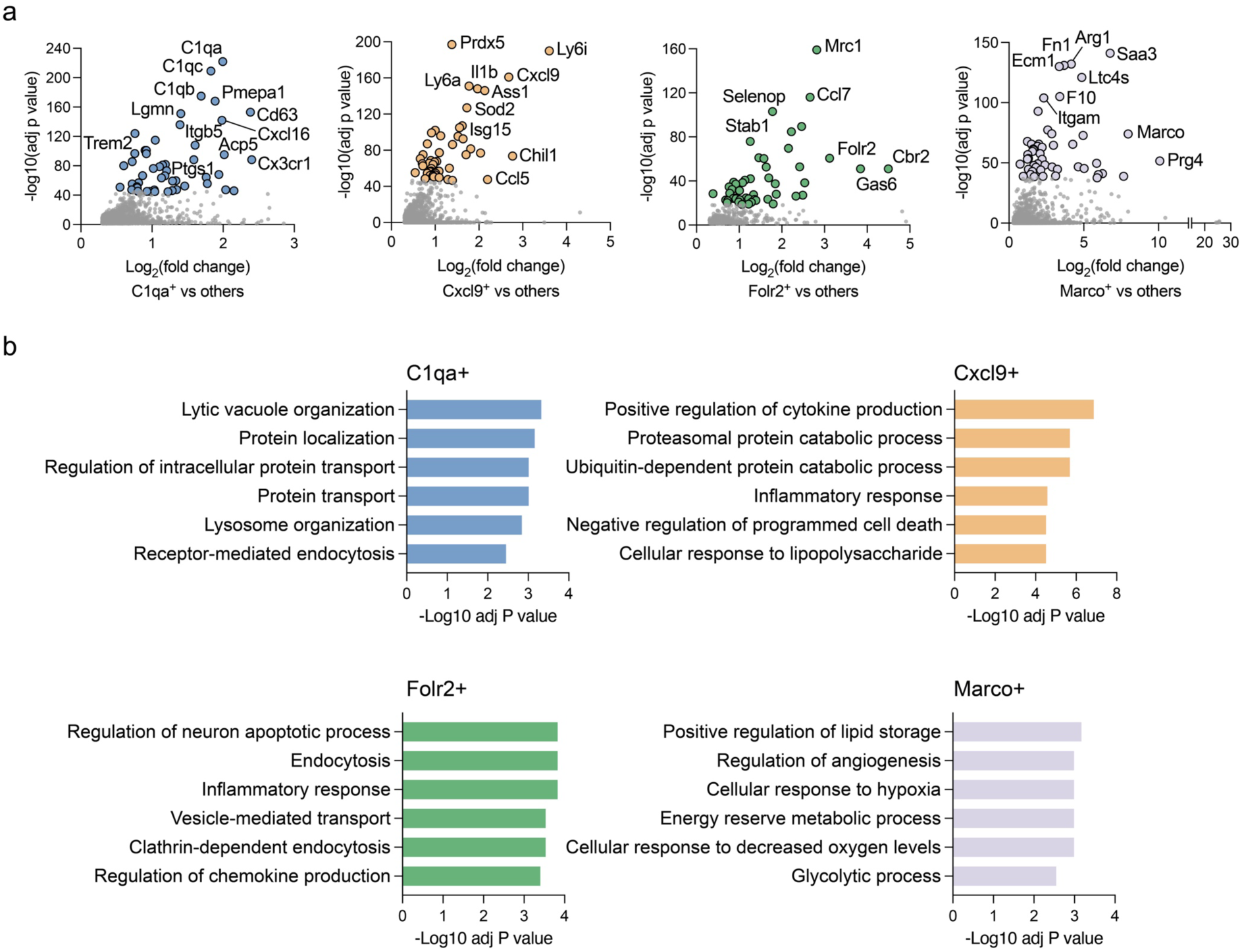
Transcriptional features of the identified tumor associated macrophage populations. **a**, Selected macrophage populations identified in the scRNA-seq experiment across control, *Serpinb2* KO, and *Serpine1* KO tumors. Top 50 genes, ordered by t-test score, are highlighted with corresponding colors in C1qa+, Cxcl9+. Folr2+ and Marco+ populations. **b**, Gene set enrichment analysis for the macrophage populations showing a log2 fold change relative to the control > 1 or <-1 upon *Serpinb2* or *Serpine1* KO. The top 6 gene ontology (GO) “biological process” gene sets enriched in each macrophage population are shown. scRNA-seq: single cell RNA sequencing, KO: knock-out.

**Figure S11.**
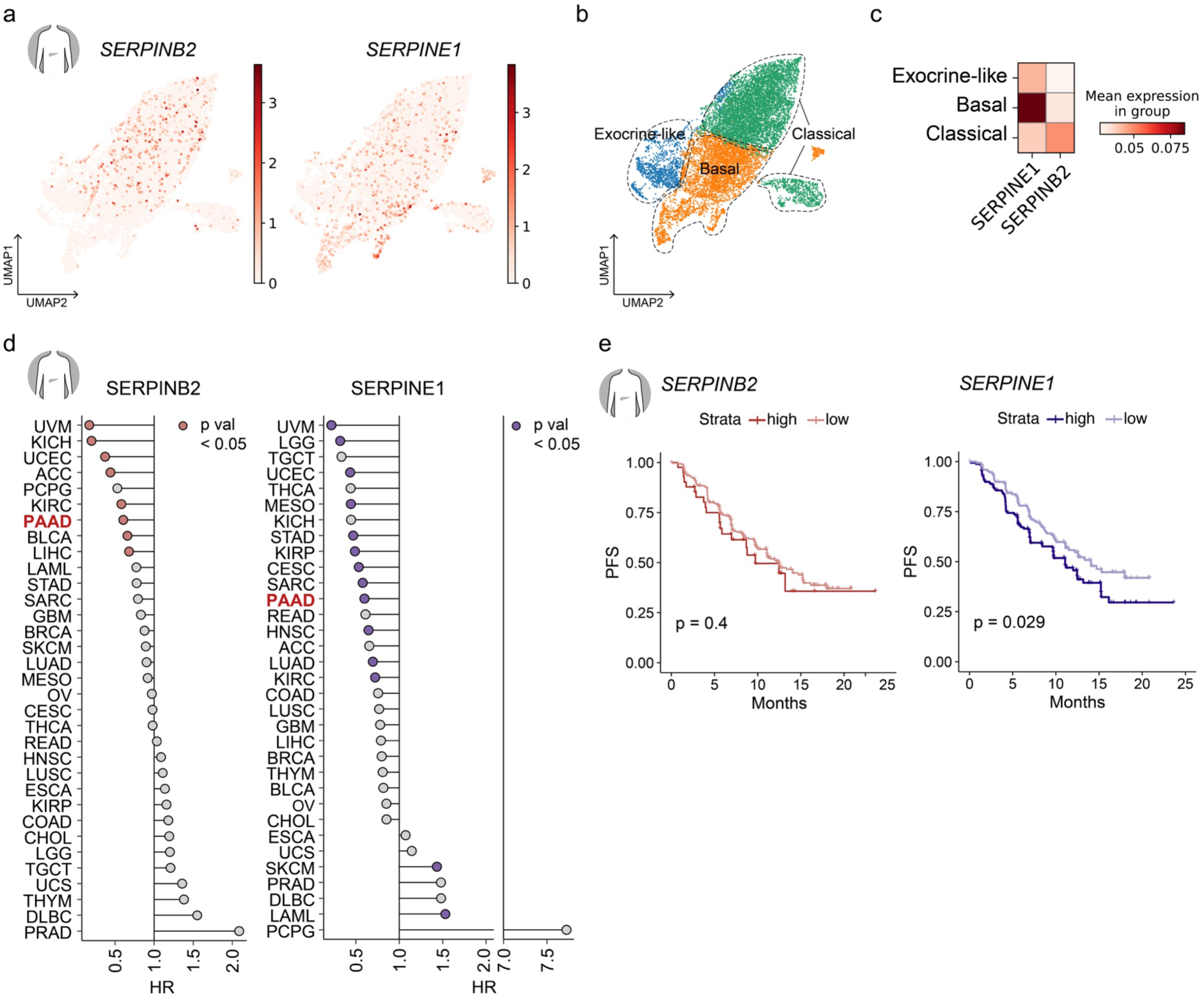
*SERPINB2* and *SERPINE1* expression characterizes tumor cells of distinct PDAC subtypes and is associated with immune checkpoint inhibitor resistance in renal cancer. **a**, UMAP projection of the tumor cell cluster from the hPDAC dataset Cui Zhou, et al.^34^, highlighting *SERPINB2* and *SERPINE1* expression. **b**, UMAP projection shown in (**a**) highlighting the annotated transcriptional subtypes from Collisson, et al.^35,36^. **c**, Heatmap displaying the mean expression of *SERPINE1* and *SERPINB2* across tumor cell transcriptional subtypes analyzed in (**b**). **d**, *SERPINB2* and *SERPINE1* gene expression is associated with worse patient overall survival in a subset of solid human cancers. Each data set within the TCGA Pan Cancer dataset was scored depending on their expression. Lollipop plots show hazard ratio (HR) scores associated with patient overall survival (OS). In color are those that were significant (P < 0.05). **e**, Kaplan–Meier survival analysis comparing progression-free survival (PFS) between patients with low versus high tumor expression of *SERPINB2* and *SERPINE1*. Stratification was based solely on the avelumab plus axitinib cohort from the JAVELIN Renal 101 trial^37^. P value in **d** was calculated by Cox proportional hazards test. The p value in **e** was calculated using the log-rank (Mantel–Cox) test. hPDAC: human PDAC, scRNA-seq: single cell RNA sequencing.

