## Supplementary table 3 for "Plasminogen activator inhibitors orchestrate the immunosuppressive tumor microenvironment in pancreatic cancer"

| REAGENT or RESOURCE | SOURCE | IDENTIFIER |
| --- | --- | --- |
| **Antibodies** | | |
| anti-AU1 tag (mouse, MICSSS, CyTOF) | Biolegend | Cat# 901901  Clone: AU1  RRID: AB_2565014 |
| anti-C tag (mouse, MICSSS, CyTOF) | Genscript | Cat# A01774  Clone: HPC4  RRID: AB_2744686 |
| anti-HA tag (mouse, MICSSS, CyTOF) | Cell Signaling | Cat# 2367S  Clone: 6E2  RRID: AB_10691311 |
| anti-HSV tag (mouse, MICSSS, CyTOF) | Thermo Fisher Scientific | Cat# PA1-26555  RRID: AB_794527v |
| anti-NWS tag (mouse, MICSSS, CyTOF) | Genscript | Cat# A01732  Clone: 5A9F9  RRID: AB_2622218 |
| anti-S tag (mouse, MICSSS, CyTOF) | Abcam | Cat# ab19321  RRID: AB_777789 |
| anti-V5 tag (mouse, MICSSS, CyTOF) | Thermo Fisher Scientific | Cat# R960-25  Clone: SV5-Pk1  RRID: AB_2556564 |
| anti-PAI1 (mouse, human; MICSSS, WB) | Invitrogen | Cat# MA1-40224  Clone: MA-33H1F7  RRID: AB_2186871 |
| anti-PAI2 (MICSSS, human; WB) | Invitrogen | Cat# PA5-27857  RRID: AB_2545333 |
| anti-Vinculin (mouse; WB) | Sigma-Aldrich | Cat# V4505  Clone: VIN-11-5  RRID: AB_477617 |
| anti-rabbit-HRP (mouse; WB) | Cell Signaling | Cat# 7074S  RRID: AB_2099233 |
| anti-mouse-HRP (mouse; WB) | Cell Signaling | Cat# 7076P2  RRID: AB_330924 |
| anti-CD4 (mouse; MICSSS) | Abcam | Cat# ab183685  Clone: EPR19514  RRID: AB_2686917 |
| anti-CD8a (mouse; MICSSS) | Cell Signaling | Cat# 98941S  Clone: D4W2Z  RRID: AB_2756376 |
| anti-CD11c (mouse; MICSSS) | Cell Signaling | Cat# 97585  Clone: D1V9Y  RRID: AB_2800282 |
| anti-F4/80 (mouse; MICSSS) | Cell Signaling | Cat# 70076  Clone: D2S9R  RRID: AB_2799771 |
| anti-FOXP3 (mouse; MICSSS) | Cell Signaling | Cat# 12653  Clone: D6O8R  RRID: AB_2797979 |
| anti-CK19 (mouse, human; MICSSS) | Abcam | Cat# ab52625  Clone: EP1580Y  RRID: AB_2281020 |
| anti-PDPN (mouse; MICSSS) | Thermo Fisher Scientific | Cat# MA5-29742  Clone:066  RRID: AB_2785565 |
| anti-aSMA (mouse; MICSSS) | Abcam | Cat# ab5694  RRID: AB_2223021 |
| anti-GZMB (mouse; MICSSS) | R&D Systems | Cat# AF1865  RRID: AB_2294988 |
| anti-Fibrin(ogen) (mouse, human; MICSSS) | Invitrogen | Cat# MA5-15906  Clone: 4H9  RRID: AB_11153062 |
| anti-LY6G (mouse; MICSSS) | Biolegend | Cat# 127601  Clone: 1A8  RRID: AB_1089180 |
| CD68 (human; MICSSS) | Invitrogen | Cat# PA5-78996  RRID: AB_2746112 |
| CD66b (human; MICSSS) | BD Biosciences | Cat# 555723  Clone:G10F5  RRID: AB_396066 |
| CD11b (mouse, human; MICSSS) | Abcam | Cat# ab133357  Clone: EPR1344  RRID: AB_2650514 |
| CD11b (FITC; mouse, flow cytometry) | BioLegend | Cat# 101206  Clone: M1/70  RRID: AB_312788 |
| CD3e (PerCPCy5.5; mouse, flow cytometry) | Invitrogen | Cat# 45-0031-82  Clone: 145-2C11  RRID: AB_1107000 |
| CD64 (APC; mouse, flow cytometry) | Invitrogen | Cat# 17-0641-82  Clone: X54-5/7.1  RRID: AB_2735010 |
| LY6G (Alexa700; mouse, flow cytometry) | BioLegend | Cat# 127622  Clone: 1A8  RRID: AB_10643269 |
| CD8a (BV510; mouse, flow cytometry) | BioLegend | Cat# 100752  Clone: 53-6.7  RRID: AB_2563057 |
| CD45 (BV650; mouse, flow cytometry) | BioLegend | Cat# 103151  Clone: 30-F11  RRID: AB_2565884 |
| **Bacterial and virus strains** | | |
| TOP10 | Thermo Fisher Scientific | Cat# C404003 |
| NEB stable competent cells | New England BioLabs | Cat# C3040I |
| **Biological samples** | | |
| Human PDAC tissue samples | Mount Sinai Tissue Biorepository | NA |
| **Chemicals, peptides, and recombinant proteins** | | |
| Puromycin | Thermo Fisher Scientific | Cat# A1113803 |
| Polybrene | Millipore | Cat# TR-1003-G |
| Bovine Serum Albumin (BSA) | Thermo Fisher Scientific | Cat# BP9705-100 |
| Cell-ID Intercalator-103 Rh | Fluidigm | Cat# 201103A |
| Cell-ID Intercalator-191/193 Ir | Fluidigm | Cat# 201192A |
| Target Retrieval Solution, pH 9 (10X) | Agilent | Cat# S2367 |
| Target Retrieval Solution Citrate, pH 6 (10X) | Aligent | Cat# S2369 |
| Protein Block, Serum Free | Agilent | Cat# X0909 |
| Hematoxylin Solution, Harris Modified | Sigma Aldrich | Cat# HHS32 |
| Antibody Diluent, Background Reducing | Agilent | Cat# S3022 |
| AEC Substrate Kit, Peroxidase (HRP) | Vector Laboratories | Cat# SK-4200 |
| Glycergel Mounting Medium | Agilent | Cat# C0563 |
| AffiniPure Fab Fragment Donkey Anti-Mouse IgG (H+L) | Jackson ImmunoResearch | Cat# 715-007-003 |
| AffiniPure Fab Fragment Donkey Anti-Rabbit IgG (H+L) | Jackson ImmunoResearch | Cat# 711-007-003 |
| AffiniPure Fab Fragment Donkey Anti-Rat IgG (H+L) | Jackson ImmunoResearch | Cat# 712-007-003 |
| AffiniPure Fab Fragment Donkey Anti-Goat IgG (H+L) | Jackson ImmunoResearch | Cat# 705-007-003 |
| EnVision+ System-HRP Labelled Polymer Anti-mouse | Agilent | Cat# 2023-04-30 |
| EnVision+ System-HRP Labelled Polymer Anti-rabbit | Agilent | Cat# 2023-02-28 |
| VisUCyte HRP Polymer Goat IgG Antibody | R&D | Cat# VC004-025 |
| ImmPRESS HRP Goat Anti-Rat IgG | Vector Laboratories | Cat# MP-7444 |
| InVivoMAb rat IgG2a | Bioxcell | Cat# BE0089 |
| InVivoMAb anti-mouse PD1 (CD279) Clone 29F.1A12 | Bioxcell | Cat# BE0273 |
| InVivoMAb rat IgG1 isotype control | Bioxcell | Cat# BE020 |
| InVivoMAb anti-CSF1 | Bioxcell | Cat# BE0204 |
| Control liposomes | Encapsula | Cat# CLD-8914 |
| Clodronate liposomes | Encapsula | Cat# CLD-8914 |
| **Critical commercial assays** | | |
| MaxPar X8 Polymer Kit | Fluidigm | Custom made |
| Foxp3 / Transcription Factor Fixation/Permeabilization Concentrate and Diluent | Thermo Fisher Scientific | Cat# 00-5521-00 |
| QIAquick PCR purification kit | Qiagen | Cat# 28104 |
| EndoFree Plasmid Maxi Kit | Qiagen | Cat# 12362 |
| ZR Plasmid Miniprep Classic kit | Zymo Research | Cat# D4015 |
| Trichrome Stain Kit (Connective Tissue Stain | Abcam | Cat# ab150686 |
| Tumor dissociation kit | Miltenyi | Cat# 130-096-730 |
| **Deposited data** | | |
| Gene Expression Omnibus | NCBI | GSE289138 |
| **Experimental models: Cell lines** | | |
| Human: 293T | ATCC | ATCC#: CRL-3216 |
| Mouse: KPC FC1245 | Tuveson laboratory | FC1245 |
| **Experimental models: Organisms/strains** | | |
| Mouse: Rag2-/- | Jackson Laboratories | Stock# 008449 |
| Mouse: H11Cas9 | Jackson Laboratories | Stock# 028239 |
| **Oligonucleotides** | | |
| See Table S1 for sequences | This paper | N/A |
| **Recombinant DNA** | | |
| Plasmid: Pro-Code vector kit | This paper/Addgene | Kit #1000000177 |
| **Software and algorithms** | | |
| Prism (v10) | Graphpad | https://graphpad.com |
| ImageJ (Fiji) (v 1.0) | Schindelin et al., 2012 | https://imagej.net/software/fiji/ |
| QuPath (v0.5.1) | Bankhead et al., 2017 | https://qupath.github.io/ |
| FlowJo (v10) | FlowJo, LLC | https://www.flowjo.com |
| Cytobank | Kotecha et al. | https://www.cytobank.org |
| Custom image processing | This paper | https://github.com/BDBrownLab |
| R (v4.2.2) |  | https://www.r-project.org/ |
| Python (v3.9) |  | https://www.python.org/ |
| Single Cell Debarcoder | Zunder et al., 2015 | https://github.com/nolanlab/single-cell-debarcoder |
| CellRanger (v5.0.1) | 10x Genomics | https://www.10xgenomics.com/support/software/cell-ranger/latest |
| Scanpy (v1.9.4) | Wolf et al., 2018 | https://squidpy.readthedocs.io/en/stable/ |
| Squidpy (v1.3.0) | Palla et al., 2022 | https://squidpy.readthedocs.io/en/stable/ |
