## Supplementary figures and images for "Plasminogen activator inhibitors orchestrate the immunosuppressive tumor microenvironment in pancreatic cancer"

### Extended figure 1

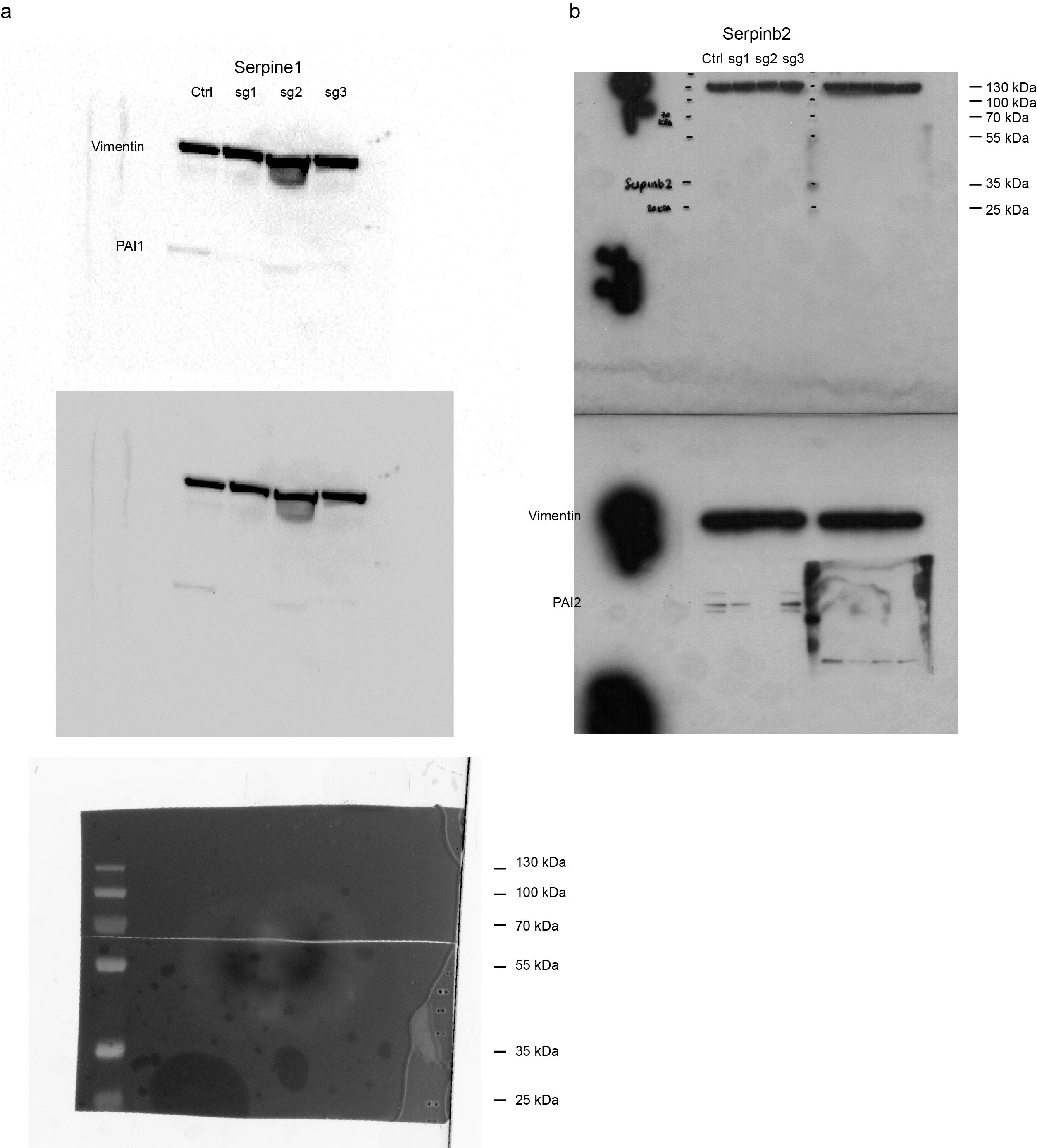

### Extended figure 2

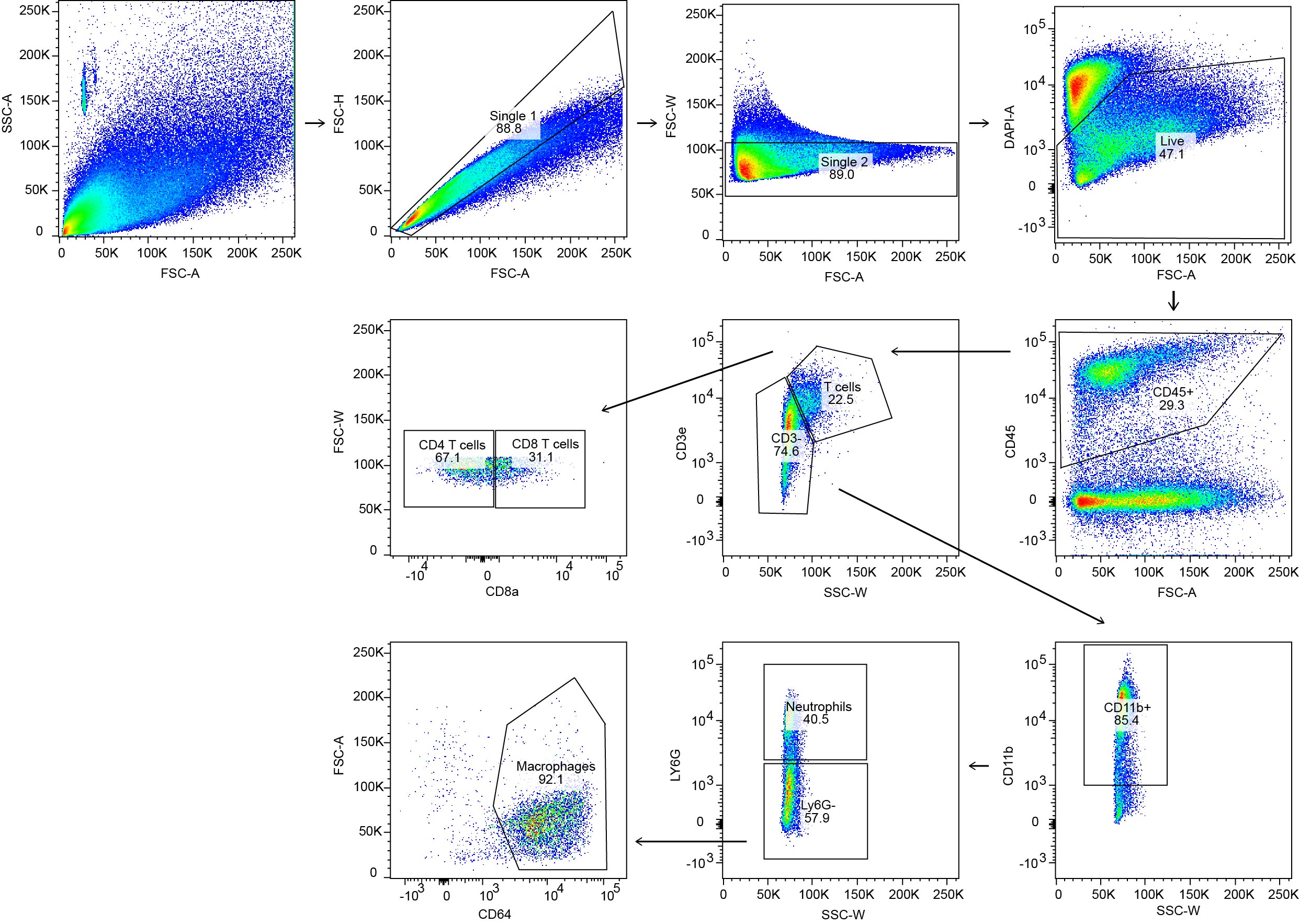
